# The integrated LIM-peptidase domain of the CSA1/CHS3 paired immune receptor detects changes in DA1 family peptidase inhibitors to confer *Albugo candida* resistance in Arabidopsis

**DOI:** 10.1101/2022.06.21.497003

**Authors:** Benguo Gu, Toby Parkes, Caroline Smith, Fu-Hao Lu, Neil McKenzie, Hui Dong, Jonathan D. G. Jones, Volkan Cevik, Michael W. Bevan

**Affiliations:** Department of Cell and Developmental Biology, John Innes Centre, Norwich Research Park, Norwich NR4 7UH, United Kingdom; The Sainsbury Laboratory, University of East Anglia, Norwich Research Park, Colney Lane, Norwich, NR4 7UH, United Kingdom; The Milner Centre for Evolution, Department of Biology and Biochemistry, University of Bath, Bath, BA2 7AY, United Kingdom

**Keywords:** . *Arabidopsis/Albugo candida*/resistance gene/LIM-Peptidase/integrated decoy/DA1 family

## Abstract

White blister rust, caused by the oomycete *Albugo candida*, is a widespread disease of Brassica crops. The Arabidopsis CSA1/DAR4 (also known as CSA1/CHS3) paired immune receptor carries an Integrated Domain (ID) with homology to the DA1 family of peptidases. Using domain swaps with DA1 family members, we show that the DAR4 ID acts as an integrated decoy for DAR3, which interacts with and inhibits the peptidase activities of DA1, DAR1 and DAR2 family members. *Albugo* infection rapidly lowered DAR3 levels and activates DA1 peptidase activity. This promotes endoreduplication of host tissues to support pathogen growth. We propose that DAR4/CSA1 senses the actions of a putative *Albugo* effector that reduces DAR3 levels and initiates defense.

## Introduction

White blister rust, caused by the oomycete *Albugo candida*, is a widespread disease of Brassica crops such as *Brassica juncea* (oilseed mustard) and *B. oleracea* vegetables [1]. White rust resistance (*WRR)* genes encoding nucleotide-binding leucine-rich repeat (NLR) immune receptors were mapped in or cloned from oilseed mustard [2, 3] and Arabidopsis [4, 5]. Most *Arabidopsis thaliana* accessions are resistant to Brassica-infecting *Albugo candida* races due to multiple NLR immune receptors [6]. However, genetic analyses of *A. thaliana* multiparent advanced generation inter-cross (MAGIC) lines identified susceptible lines that enabled cloning of multiple *WRR* genes against Brassica infecting *A. candida* races. [6]. This repertoire of NLR genes was proposed to recognise a range of *A. candida* race-specific effectors [5].

Genetic mapping of Col-5- and Ws-2- derived Arabidopsis Recombinant Inbred Lines (RILs) identified a further three loci that confer resistance to *A. candida* AcEM2 [4, 5]. Fine mapping of the *WRR5* locus identified two resistance genes, *WRR5A* (At5G17880) and *WRR5B* (At5G17890) in a head-to-head configuration. Both *WRR5A* and *WRR5B* were required for conferring chlorotic resistance in transgenic lines of the susceptible accession Ws-2 to *A. candida* AcEM2 [4]. Previously these genes had been identified as *CSA1* (*WRR5A*) and *CHS3* (*WRR5B*) [7]. The *chs3-2D* allele caused auto-immune responses requiring *CSA1*, analogous to the requirement of RPS4 for the autoimmunity conferred by RRS1^slh1^, suggesting CSA1 and CHS3 also form a paired immune receptor [8]. The *chs3-2D* autoimmune allele carries a point mutation in the LIM (Lin-11, Isl-1, and Mec-3)-peptidase integrated domain (ID) of CHS3/WRR5B, and this ID is also found in multiple genomes in the Brassicaceae, Fabaceae and Rosaceae families [9]; [10].

Recent analyses of natural variation of *CSA1/CHS3* in *Arabidopsis thaliana* identified three clades, with clade I carrying an ID that maintains the CSA1/CHS3 complex in an inactive state. Members of all three clades interact with the BAK1-BIR1 brassinosteroid signalling complex that maintains their inactivity [5, 11]. Interestingly, clade I complexes maintain this function while gaining an additional function from its ID. This function remains uncharacterised.

IDs have been shown to perceive pathogen effectors and their biochemical activities, likely directly. Analyses of ID functions show they are homologous to proteins that are pathogen effector targets. When integrated into NLRs, they enable recognition of effector activities and activation of defense [8,12–14]. They have been dubbed “integrated decoys” as they detect effector activities on authentic virulence targets [10,12,15]. Effector molecules produced by pathogens facilitate colonisation and reproduction in plant hosts by subverting diverse host cell functions and suppressing innate resistance [16, 17]. *Albugo* is an obligate biotroph that infects leaf cells by invaginating the plasma membrane and forming haustoria, which then take up nutrients produced by living host cells. Multiple aspects of host plant growth and metabolism are modified by effectors to support pathogen biotrophic growth, and increasing knowledge of the diverse mechanisms by which effectors modulate plant processes are generating new insights into both the regulation of normal host growth and of infection processes [18, 19]. Despite exciting progress in understanding ID-effector interactions, most, including the widespread LIM-peptidase IDs, remain poorly understood.

The LIM- peptidase protein domains characterise the eight members of the *DA1* family of growth regulators in Arabidopsis Col-0 [20]. The LIM domain of DA1 comprises two pairs of Zn-finger motifs in a characteristic conformation that confer specificity to protein-protein interactions [21], and a conserved C-terminal Zn metallopeptidase domain with an (A)HEMMH(A) active site [21, 22]. The DA1, DAR1 and DAR2 members of the DA1 peptidase family proteolytically cleave and inactivate, *via* N-end rule mediated proteolysis, diverse proteins involved in cell proliferation to modulate organ size by limiting cell numbers during organogenesis [20, 22]. DA1 peptidases also regulate auxin responses by cleaving Transmembrane Kinase I to release an intracellular kinase domain that relocates from the plasma membrane to the nucleus to suppress auxin responses [23]. *DAR4* is *CHS3/WRR5B*, carrying TIR, LRR and NB-ARC domains typically found in NLR resistance genes [20], and *DAR5* carries RPW8 and NB-ARC domains. This suggests that the activities of DA1 family members may be modified by pathogen effectors to support pathogen growth, and that these immune receptors have evolved to detect this activity.

Gaining a comprehensive understanding of the interplay between pathogens and hosts is a key objective as it provides both new insights into the regulation of host plant growth and a means of predicting pathogenicity and host resistance responses based on genetic and genomic analyses of effector and *R* gene repertoires [24, 25]. Using domain swaps, we show that the LIM-peptidase domain of DAR4/CHS3/WRR5B is an integrated decoy for the DAR3 family member. *Albugo candida* race AcEM2 induces endoreduplication in infected leaf tissues and this requires DA1 and DAR1 function, defining them as susceptibility genes. DA1 family peptidase activities are reduced by formation of complexes with other family members [26], including DAR3 and DAR7. AcEM2 infection led to the rapid degradation of DAR3, activating DA1 peptidase activity, thereby promoting endoreduplication and pathogen growth.

## Results

### The LIM-Peptidase (LP) domain of DAR4 is a target for an *Albugo* effector

*DAR4/CHS3/WRR5B* and *CSA1* genes are in head-to-head configuration on chromosome 5 in *Arabidopsis thaliana* Col-0 [7]. To verify that they encode proteins that depend on each other for immune receptor function, the *chs3-1* and *chs3-2d* mutants that confer autoimmunity [27] were transiently co-expressed with *CSA1* in *Nicotiana tabacum*. Both *chs3* mutants triggered hypersensitive responses (HR) and strong cell death reactions in the presence but not the absence of CSA1. Wild type DAR4 co-expressed with CSA1 did not trigger HR (Figure 1, Supplementary Figures 1, 4A and 4B). To confirm their mutual dependence, DAR4 and CSA1 constructs were co-expressed with another R protein pair, RRS1 and RPS4 [9], and tested for HR in *N. tabacum*. As expected, no HR was triggered by the non-cognate protein pairs (Supplementary Figure 2 and 3). These observations confirm that DAR4/CHS3 and CSA1 form a functional R protein pair [5].

**Figure 1.**
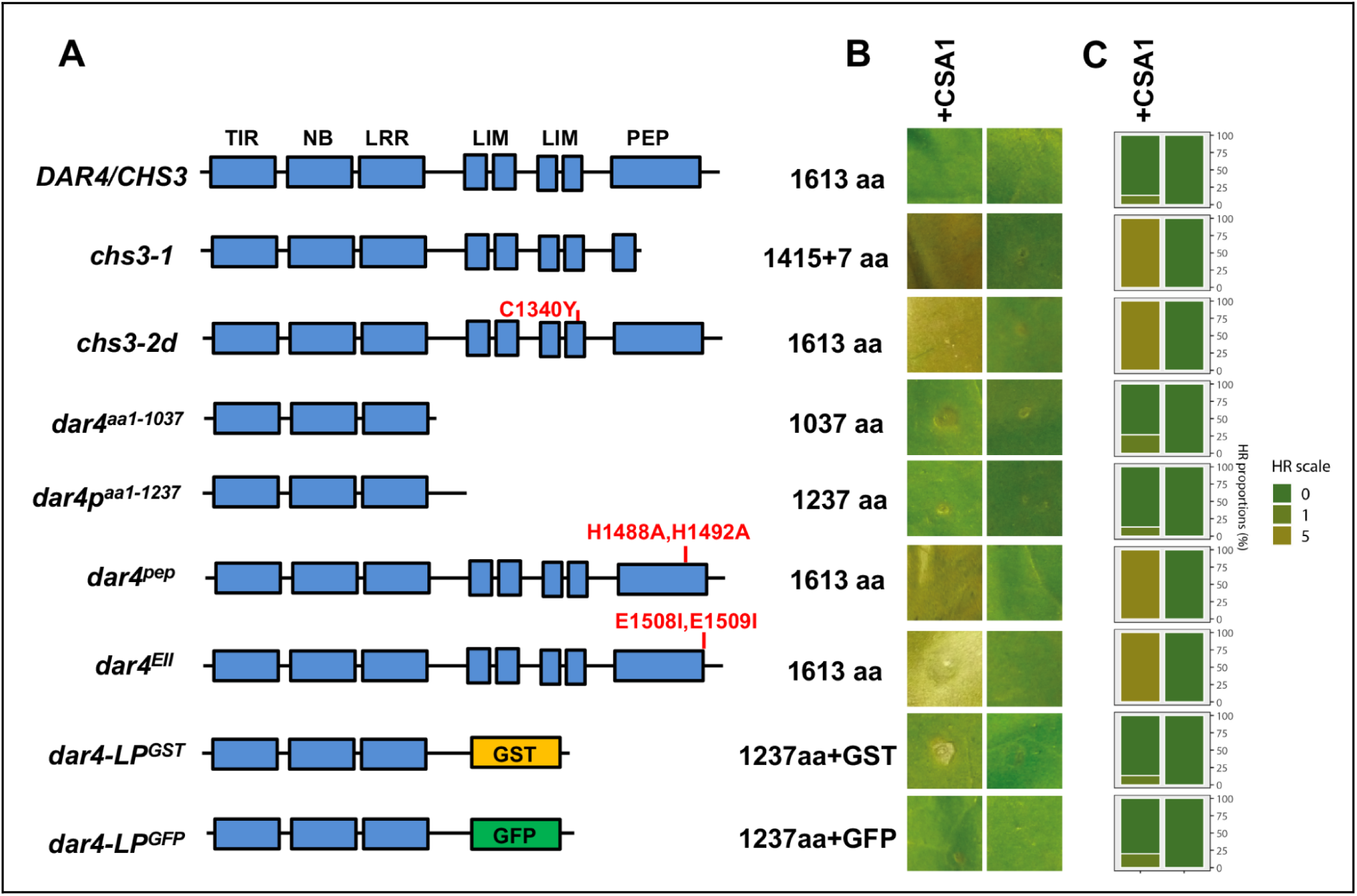
The LIM-peptidase region of DAR4 functions as an integrated decoy. (**A**) Diagram of known protein motifs of DAR4 and mutant and chimaeric forms used to test hypersensitive responses. TIR: Toll and Interleukin-1 Receptor homology region; NB: NB-ARC Nucleotide-Binding ARC,APAF-1,R proteins and CED-4 domains; LRR: Leucine-Rich-Repeat domain; LIM: LIN11, Isl1, MEC3 domains; Each Zn finger of the two LIM domains is represented; PEP: Zinc-metallopeptidase region with AHEMMHA–-EE active site. GST: Glutathione S-transferase; GFP: Green Fluorescent Protein. Red letters indicate the positions of mutated amino acid codons. C1340Y is in the fourth Zn-finger of the LIM repeats. H1488, H1492, E 1508, E1509 are in the AHEMMA and EE motifs of the peptidase active site. The diagram is not to scale. Supplementary Figure 5 shows the amino acid sequences and motifs of the DA1 family. (**B**) Hypersensitive responses in *N. tabacum* leaves were assessed at 4 days post infection. Images represent typical samples of three replicates. DAR4 and variants in panel A were co-expressed with (+) or without CSA1. The expression levels of DAR4 variants is shown in Supplementary Figure 4. (**C**) Quantified HR responses. The stacked bars are colour coded to show the percentage of each cell death scale (0-5) in 15 panels for each sample. Yellow represents the presence of HR, green represents the absence of HR [28].

The *chs3-1* allele truncates the C-terminal region containing the peptidase active site and the *chs3-2D* allele disrupts the 4th Zn finger of the LIM domain of DAR4 [20, 27] (Figure 1). This disruption in DA1 (through the equivalent C274Y mutation) leads to loss of DA1 peptidase activity [22]. To test whether the integrity of the peptidase active site in DAR4 is required for its regulated immune receptor function, two active site mutations that abolish DA1 peptidase activity were introduced into DAR4: H1488A and H1492A ( *dar4^pep^*) and E1508I and E1509I (*dar4^EEII^*) [22]. These mutant DAR4 proteins were co-expressed with CSA1 in *N. tabacum.* All of these mutations (the *chs3-1* truncation, *chs3-2D* Zn-finger disruption and the two peptidase active site mutants) trigger HR in a *CSA1*-dependent way (Figure 1, Supplementary Figures 4A and 4B), indicating that a functionally intact LIM-Peptidase domain in DAR4 is required to maintain the capacity of DAR4/CHS3 to deliver appropriately regulated CSA1-dependent defense responses, while DAR4/CHS3 carrying mutated LIM-Peptidase domains trigger CSA1-dependent HR. Thus, the LIM-Peptidase domain of DAR4 acts as a potential integrated decoy. To test this further, *dar4^aa1-1037^* and *dar4^aa1-1237^* premature termination truncations of the LIM-Peptidase domain were made, and two chimeric proteins that swapped the LIM-Peptidase domain with Green Fluorescent Protein (*dar4-LP^GFP^* or Glutathione S-Transferase (*dar4-LP^GST^*) were also made and co-expressed with CSA1 in *N. tabacum* leaves. None of these deletions or domain swaps triggered HR (Figure 1, Supplementary Figures 4A and 4B). The triggering of HR by *chs3-1*, *chs3-2d*, *dar4^pep^* and *dar4^EEII^*, and the absence of HR triggered by DAR4 premature termination variants without a LIM-Peptidase domain, indicates that the LIM-Peptidase domain regulates the immune functions of the R protein pair, DAR4 and CSA1. In its native conformation it suppresses immune responses, while changes to the functional integrity of the LIM-Peptidase domain trigger CSA1-dependent HR.

Analyses of genomic data suggested that IDs arise by duplication and recombination of host gene regions that are targets of pathogen effectors [29]. For example, the WRKY ID of RRS1 and its homologous WRKY transcription factor are both acetylated by the bacterial effector PopP2 [8, 30], triggering HR through RRS1 and RPS4. Therefore, if the DAR4 LIM-peptidase domain responds to putative pathogen effectors that may target other members of the DA1 family, there should be structural and functional conservation between the DAR4 LIM-Peptidase domain and other family members targeted by putative effectors. The protein sequence relationships and domains of the eight DA1 family members are shown in Figure 2A. DAR3 is the family member most closely related to DAR4. DAR3 and DAR7 are characterised by the absence of Ubiquitin Interaction Motifs (UIMs) and may not be functional DA1 family peptidases [22]. To test the functional conservation of the DA1 family LIM-Peptidase domains in triggering HR responses, the LIM-Peptidase domains of the seven other members of the DA1 family were fused with the NLR region of DAR4 (Figure 2B, Supplementary Figure 4A, 4C and Table 2). These were co-expressed with CSA1 to test their ability to trigger HR in *N. tabacum*. dar4-LP (DA1, DAR1, DAR2, DAR3 and DAR7) did not trigger HR, with phenotypes that were the same as wild type DAR4. In contrast, dar4-LP(DAR5) and dar4-LP(DAR6) fusions triggered HR (Figure 2B, Supplementary Figures 4A and 4C). As shown in Figure 1, fusions of the DAR4 NLR region with the un-related proteins, dar4-LP^GST^ and dar4-LP^GFP^, also did not trigger HR. To exclude the possibility that swapped DA1 family LIM-Peptidase domains cannot generate an HR signal, a mutant of dar4-LP(DA1) called dar4-LP(DA1)-2D with an amino acid alteration (DA1 C274Y) corresponding to the mutation in *chs3-2D* was tested. Strong HR was triggered by co-expressing this protein and CSA1 (Figure 2B, Supplementary Figures 4A and 4C), demonstrating the capacity of fusions of DA1 family LIM-Peptidase domains to trigger HR. As DA1 is more distantly related to DAR4 compared to DAR3 and DAR7 (Figure 2A and Supplementary Figure 5), it is likely that the LIM-Peptidase domains of DA1, DAR1, DAR2, DAR3 and DAR7 all share a functionally conserved potential with DAR4 to trigger HR when fused to the DAR4 NLR region. Therefore, DAR4 is probably an R protein for these family members. As DAR5 and DAR6 LIM-peptidase fusions trigger HR when fused to DAR4, this suggests they are structurally and functionally different from other family members (Figure 2).

**Figure 2.**
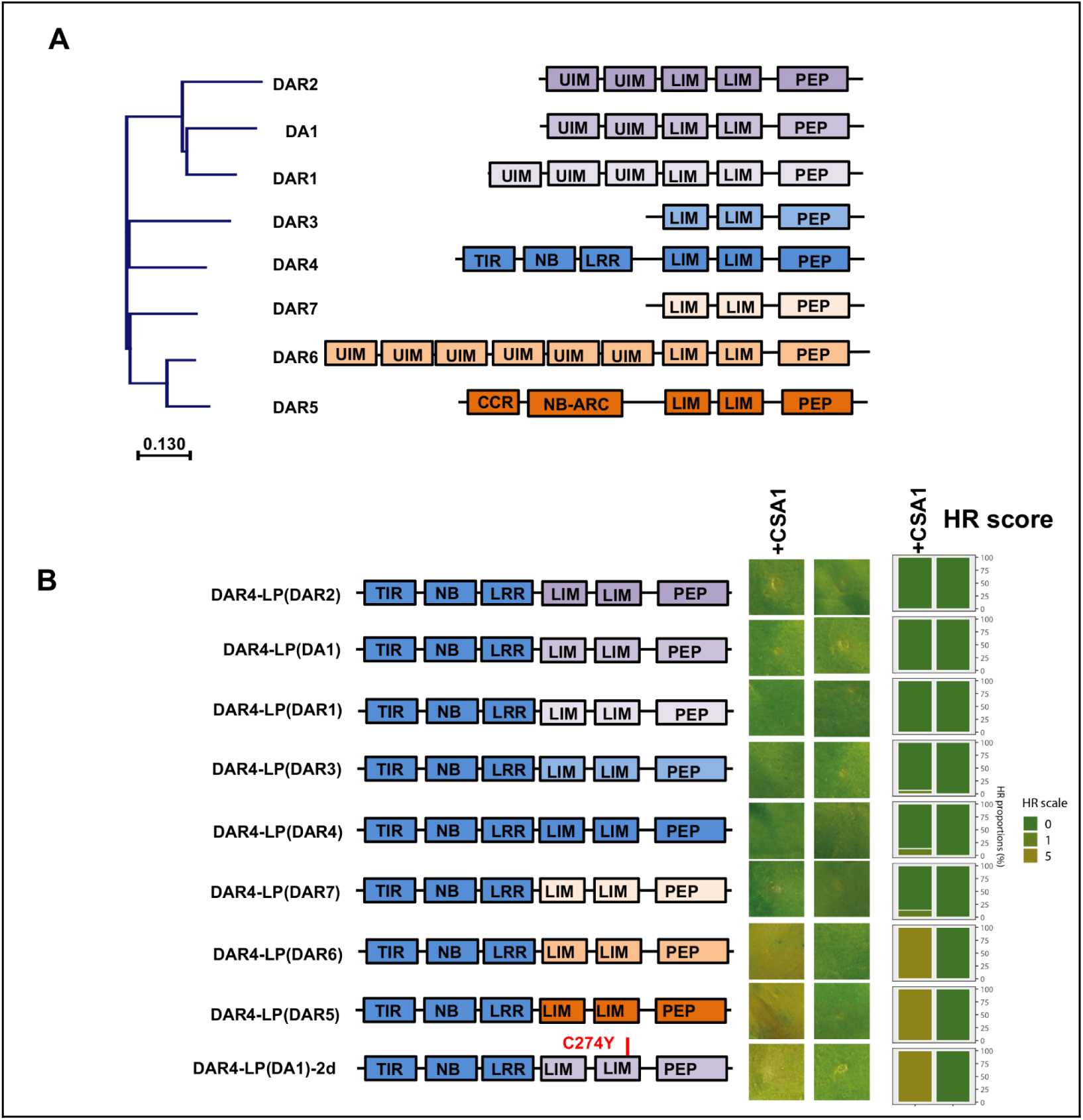
Domain swaps of LIM-peptidase regions of DA1 family members into DAR4 elicit HR responses. (**A**) Left: a Neighbour-Joint phylogenetic tree of LIM-peptidase region amino acid sequences of the DA1 family in *Arabidopsis thaliana* Col-0. The distance scale is shown at the bottom of the panel. Right: Diagram of the structural motifs of members of the DA1 family in Arabidopsis thaliana Col_0. UIM: Ubiquitin Interaction Motif; LIM domains; PEP: Peptidase domain; TIR-NB-ABC-LRR region; CCR; Coiled-Coil of RPW8. The diagram is not to scale. (**B**) Diagram of LIM-peptidase regions of DA1 family members swapped with that of DAR4. The colours indicate the different LIM-Peptidase (LP) regions of family members in chimaeric DAR4 proteins. DAR4-LP(DA1)-2d is DAR4 containing the LIM-Peptidase region of DA1 with the DA1 C274Y amino acid change, shown in red. The right panel shows hypersensitive responses (HR) of chimaeric DAR4 proteins expressed with (+) and without CSA1. HR responses were quantified. The stacked bars are colour coded to show the percentage of each cell death scale (0-5) in 15 samples. Yellow represents the presence of HR, green represents the absence of HR. The diagram is not to scale.

### The LIM-Peptidase domain of DAR4 lacks the activity of DA1 family peptidases

DA1, DAR1 and DAR2 are functionally redundant closely-related members of the DA1 family (Figure 2A) [20], and share substrates of their peptidase activities [20, 22]. We therefore selected DA1 to represent the activities of DAR1 and DAR2 in comparison to DAR4. To test whether the LIM- Peptidase domain of DAR4 shares peptidase substrates with DA1, the LIM- Peptidase domain of DAR4 was swapped with that of DA1 to make da1-LP(DAR4) and its potential to cleave the Big Brother E3 ligase (BB-3FLAG) in *da1-ko1dar1-1* leaf protoplasts was tested. No cleavage of BB-3FLAG was detected (Figures 3B and 3C), showing functional variation between LIM- Peptidase domains of DA1 and DAR4.

**Figure 3.**
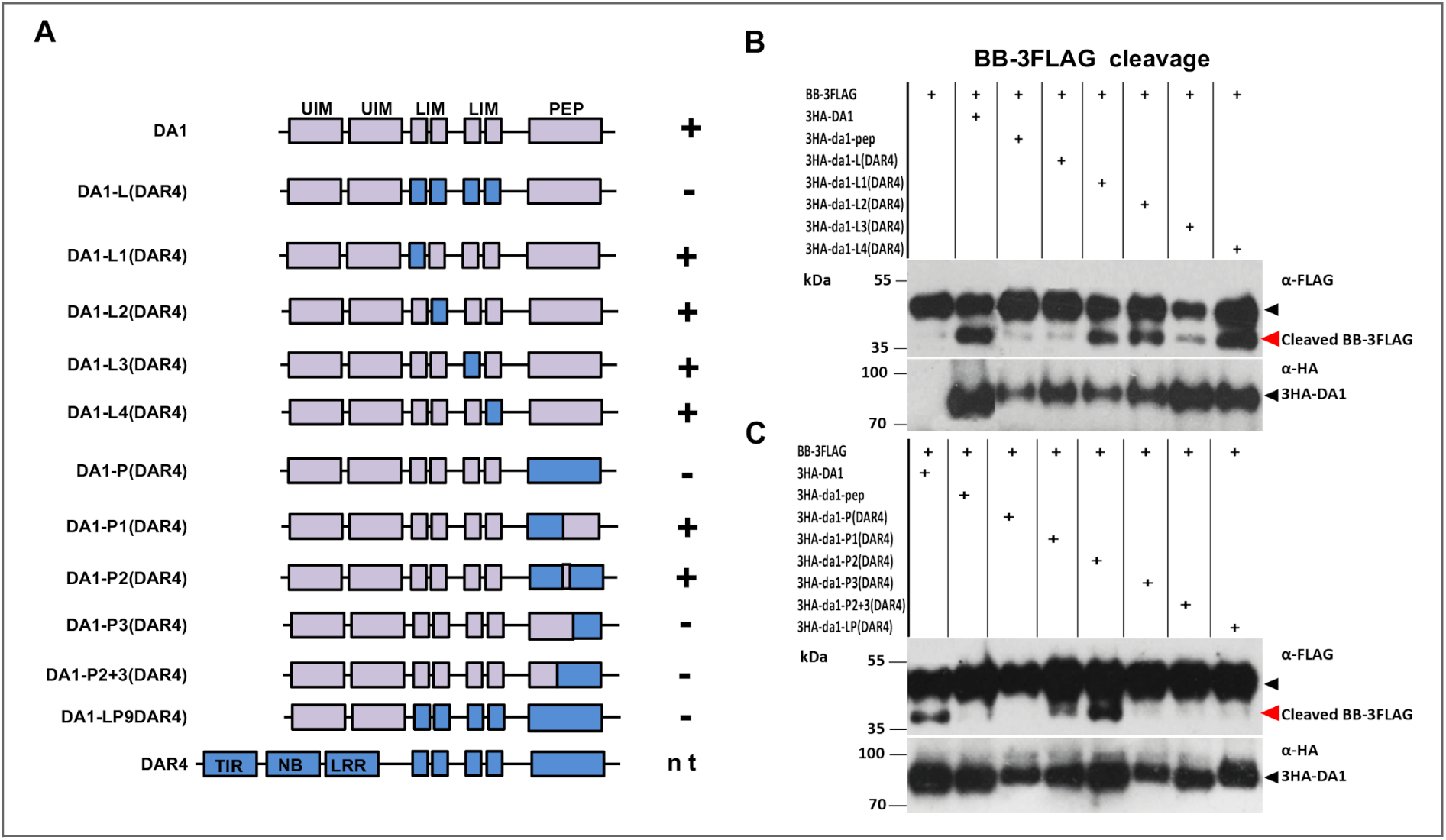
Peptidase activities of LIM-peptidase regions of DAR4 in chimaeric DA1 proteins. (A) Diagram of chimaeric 3HA-DA1 proteins used to test peptidase activities. Lilac: DA1 regions. Blue: DAR4 regions. The domain names are described in the Legend to Figure 3. The diagram is not to scale. **+** shows BB-3FLAG cleavage, **-** no cleavage, nt not tested (panels B and C). (B) and (C) Immunoblot of BB-3FLAG cleavage reactions in transfected *da1dar1* mesophyll protoplasts using 3HA-DA1 chimaeric proteins. 3HA-DA1 is a positive control and the 3HA-DA1 peptidase mutant (3HA-DA1^pep^) is a negative control. BB-3FLAG proteins were detected by anti-FLAG-HRP antibodies. 3HA-DA1 proteins were detected by anti-HA-HRP antibodies. The positions of full-length (black arrow) and cleaved (red arrow) BB-FLAG proteins are indicated.

As there was functional conservation of the LIM- Peptidase domains of DA1 and DAR4 with respect to triggering HR in *N. tabacum* (Figures 2C and 2D), smaller regions of the DAR4 LIM- Peptidase domain were swapped with that of DA1 and tested for their peptidase activity on BB-3FLAG. Neither the LIM domain nor the peptidase domain of DAR4 in DA1 could facilitate BB cleavage (Figure 3B and 3C). However, domain swaps of smaller regions, comprising each of the four single zinc fingers of the LIM and LIM-like domains, the conserved region between the LIM-like domain and the peptidase active site, and a small active site region of DAR4 all efficiently cleaved BB-3FLAG in the DA1 protein context (Figure 3B and 3C). This demonstrated that functional conservation was limited to smaller regions of the LIM Peptidase of DA1 and DAR4. A failure to ubiquitylate the DA1 domain swap proteins was ruled out as an explanation for reduced BB cleavage by the domain swap proteins as they were all ubiquitylated (Supplementary Figure 6). This functional variation between the LIM- Peptidase domains of DA1 and DAR4, and the functional conservation of DA1, DAR1 and DAR2 [31], indicated that these three family members may not be direct targets of putative effectors that are sensed by the DAR4 R protein, and that DAR4 may have other features recognised by putative effector(s). It is therefore possible that the more closely related family members (DAR3 and DAR7), which also did not trigger HR when their LIM- peptidase regions were fused to DAR4 (Figure 2B), may have a more direct role in DAR4- mediated resistance, for example as effector targets, than DA1, DAR1 and DAR2.

### *DA1* family peptidases confer susceptibility to diverse plant pathogens

The functions of DAR3 and DAR7 are not known, and possible roles of the relatively well-characterised DA1 and DAR1 family members in pathogen responses have also not been studied. We therefore assessed the involvement of *DA1* and *DAR1* in pathogen growth by measuring the growth of two model pathogens, the bacterial pathogen *Pseudomonas syringae* pathovar *tomato* (*Pst*) strain DC3000 and the oomycete pathogen *Hyaloperonospora arabidopsidis* (*Hpa*) isolate Noco2, in *da1dar1* loss of function double mutants and *DA1* over-expression lines in Col-0.

A *DA1* single loss-of-function mutant, *da1-ko1*, showed no significant differences in *Pst* DC3000 growth compared to Col-0, while a double *da1-ko1dar1-1* mutant supported much lower levels of pathogen growth. A single *DAR1* loss of function mutant, *dar1-1*, had an intermediate growth phenotype between Col-0 and the double mutant (Figure 4B and Supplementary Table 4). Similar, but more extreme, reductions in pathogen growth were observed after infection by *Hpa* isolate Noco2. The double mutant *da1-ko1dar1-1* exhibited strongly reduced Noco2 growth, while the two single mutants, *da1-ko1* and *dar1-1*, exhibited moderately reduced Noco2 growth compared with wild type Col-0 (Figure 4B and Supplementary Table 3). No significant differences in Noco2 growth were detected in the *DA1* over-expression line *ox-DA1-1.3* (Figure 4B and Supplementary Table 3). Therefore, *DA1* and *DAR1* are redundantly required for optimal growth of bacterial and oomycete pathogens in Arabidopsis, and are therefore susceptibility genes to bacterial *Pst* DC3000 and oomycete *Hpa* Noco2 pathogens.

**Figure 4.**
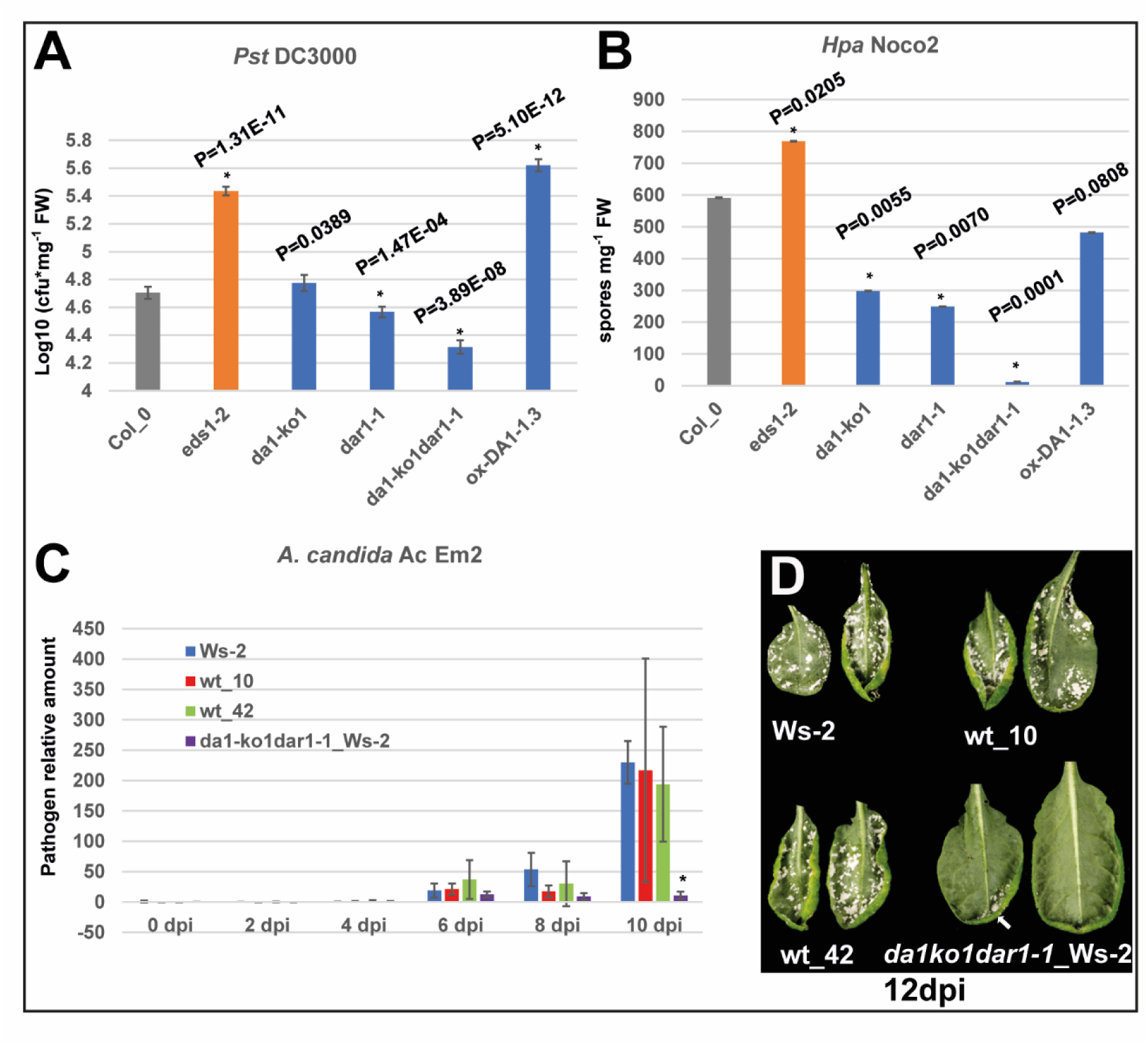
*DA1* and *DAR1* facilitate growth of bacterial and oomycete pathogens in *Arabidopsis thaliana*. (A) *Pseudomonas syringae Pst* DC3000 growth on *Arabidopsis thaliana* Col-0 *da1* and *dar1* knock-out lines and a *35S::DA1* over-expressing line. cfu: colony-forming units per mg fresh weight of inoculated leaves. The Col-0 *eds1-2* mutant was used as a standard susceptible control. (B) *Hyaloperonospora arabidopsidis* (*Hpa)* race Noco2 growth on *Arabidopsis thaliana* Col-0 *da1* and *dar1* knock-out lines and a *35S::DA1* over-expressing line. FW: fresh weight of spores in infected plants. The Col-0 *eds1-2* mutant was used as a susceptible control. C) Time-course of *Albugo candida* AcEM2 growth on *Arabidopsis thaliana* Ws-2 and backcross lines with Col-0 *da1dar1*. wt-10 and wt-44 contain wt *DA1* and *DAR1* loci from Ws-2 in BC_7_F_2_, while *da1-ko1dar1-1*_Ws-2 contains homozygous *da1dar1* mutants in BC_7_F_2_. Infection progress was measured using the ratio of pathogen:host DNA levels. dpi days post-infection. Statistical significance was at *P < 0.01 based on a two-tailed Student’s t test. Comparisons of pathogen levels were those in Ws-2 at 0 dpi. Error bars represent SD of the mean. (D) Image of *Albugo candida* AcEM2 growth on leaves of Ws-2 and backcrossed lines at 12 dpi leaves. The white areas are Em2, and the arrow indicates a small amount of pathogen growth on the *da1-ko1dar1-1_*Ws-2 backcrossed line.

As *DAR4* and *CSA1* comprise a resistance gene, *WRR5*, for *A. candida* [4] it is possible that the activities of DA1 family members may be required for AcEM2 growth, and that effector molecules detected by DAR4/CSA1 may modulate the activities of DA1 family members to support pathogen growth. To test this, the *da1-ko1dar1-1* double mutants in Col-0 were backcrossed into the *A. thaliana* Ws-2 accession, which is susceptible to *A. candida* AcEM2. In BC_7_F_2_, and a homozygous double mutant line (*da1-ko1dar1-1*_Ws-2) and two lines homozygous for the Ws-2 wild-type *DA1* and *DAR1* alleles (wt-10 and wt-42) were tested for AcEM2 growth. This was strongly reduced in the *da1-ko1dar1-1* line, while both wild type *DA1 DAR1* Ws-2 lines and the Ws-2 parental line supported similar high levels of pathogen growth (Figure 4C, 4D and Supplementary Table 5). AcEM2 growth was also assessed in a transgenic *DA1* over-expression line (*35S::DA1-GFP_*Ws-2) in Ws-2. No differences in AcEM2 growth were detected (Table 5 and Supplementary Figure 7) . These observations support the hypothesis that *DA1* and *DAR1* are required for optimal pathogen growth and are therefore susceptibility genes.

### Leaf cell endoreduplication induced by *A. candida* AcEM2 infection requires DA1 family peptidase activity

*DA1*, *DAR1* and *DAR2* promote the transition from mitosis to endoreduplication during organ growth by peptidase-mediated cleavage of positive regulators of mitosis that include TCP14, TCP15 and UPB15 ([20,22,31][32]. Endoreduplication is also induced during pathogen infection to facilitate pathogen growth [33–35]. To test if DA1 family peptidase activities contribute to pathogen-induced endoreduplication, ploidy levels of leaf cells during infection by oomycete pathogen *A. candida* AcEM2 were measured in susceptible Ws-2 and the wt-42 DA1 DAR1 Ws-2 BC7F2 wild type lines. Both lines showed similar increases in leaf cell ploidy 6 days after inoculation (dpi) (Figure 5A, 5B, Supplementary Figure 8A, 8B and Table 6).

**Figure 5.**
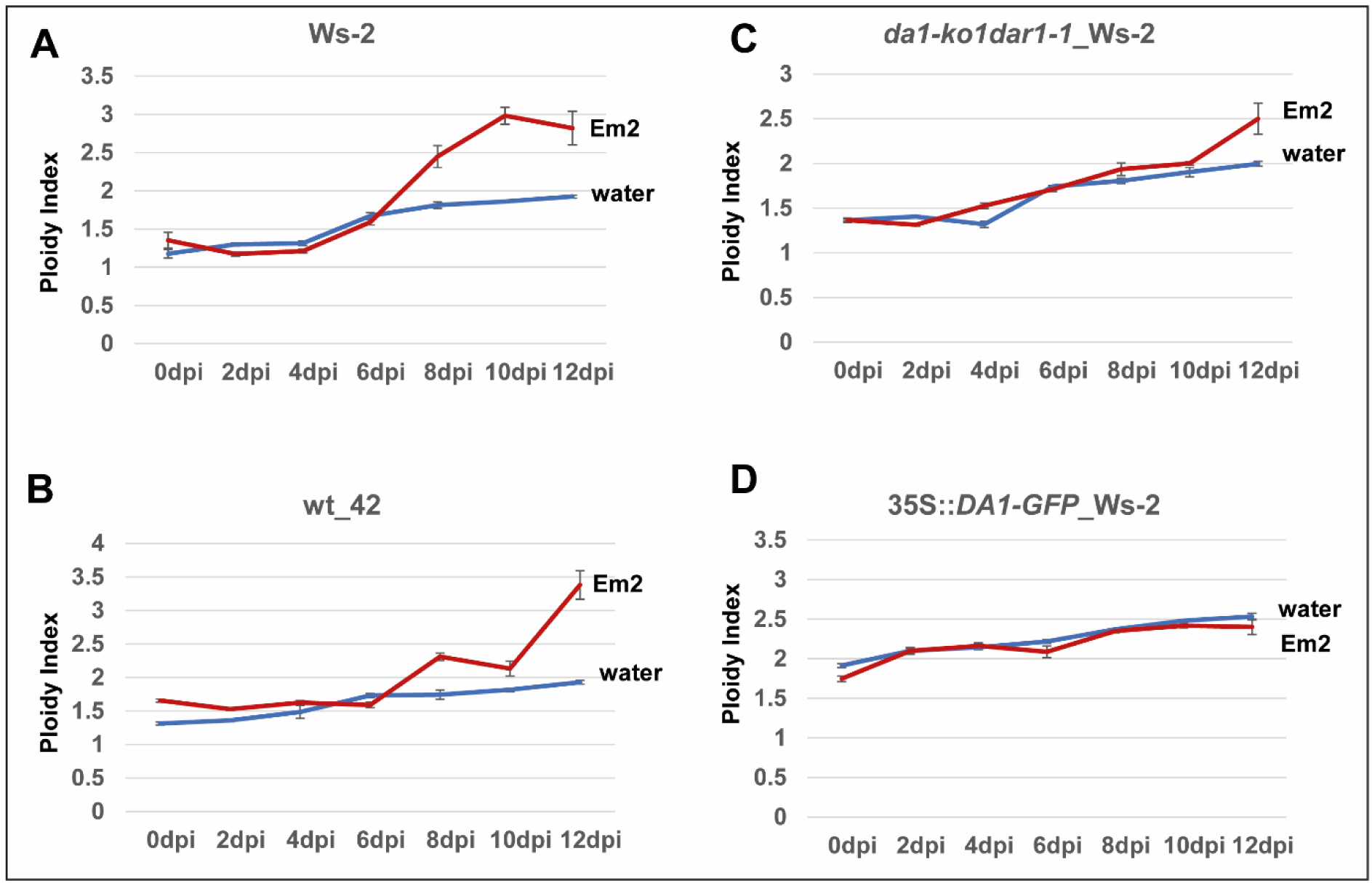
Increases in leaf ploidy levels during *A. candida* Ac Em2 infection of *Arabidopsis thaliana* are dependent on DA1 and DAR1. The graphs show ploidy levels of infected leaf cells during *A. candida* AcEM2 infection of *Arabidopsis thaliana* Ws-2 wt, wt-42, *da1-ko1dar1-1*_Ws-2 backcross line leaves, and a *35S::DA1* over-expression line in Ws-2 up to 12 days post infection. The red lines show inoculation with AcEM2 spores and the blue lines show mock inoculation with water. (A) wild-type Ws-2 ploidy index (B) Ws-2 backcross line wt-42 containing wt Ws-2 *DA1* and *DAR1* genes (C) Ws-2 backcross line containing loss-of-function Col-0 *da1-ko1* and *dar1-1* genes. (D) *35S::DA1* over-expression line. Note the increased ploidy index at 0 days post infection. Error bars represent SD of the mean.

In comparison, the *da1dar1* double mutant in Ws-2 exhibited delayed ploidy increases, with relatively small increases at 12 dpi (Figure 5C, Supplementary Figure 8C and Table 6). A *DA1* over-expression line (35S::*DA1-GFP_*Ws-2) exhibited higher ploidy levels at all stages of AcEM2 growth (Figure 5D, Supplementary Figure 8D and Table 6) compared to wild-type Ws-2 (Figure 5, Supplementary Figure 8 and Table 6 ), although AcEM2 growth was not detectably increased in this line (Table 5 and Supplementary Figure 7). This indicated that ploidy increases may influence AcEM2 growth at later stages of infection. Collectively, these data show that *A. candida* AcEM2 promotes *DA1*- and *DAR1*- dependent host cell endoreduplication and ploidy increases to support its growth.

### DAR3 and DAR7 inhibit the peptidase activities of DA1 family peptidases

Analyses of the LIM- Peptidase domains of DA1 family members demonstrated functional differences between DAR4 and DA1, DAR1 and DAR2 (Figure 3), suggesting that DAR3 and DAR7, which are more closely related to DAR4, may be potential targets of putative pathogen effectors whose effects are detected by the ID of the DAR4/CSA1 immune receptor. DA1 interaction with itself modulates its peptidase activity [26]. We showed that DAR3 also interacts with itself and DA1, DAR1 and DAR2 using co-immunoprecipitation of protoplast-expressed proteins (Figure 6A). To understand the functions of these interactions, DAR3-GFP was co-expressed with 3HA-DA1 and its substrate BB-3FLAG in Col-0 *da1dar1* mesophyll protoplasts to test its influence on BB-3FLAG cleavage by DA1. GFP was used as a co-expression standard in each protoplast transformation. BB-3FLAG cleavage was dramatically reduced by DAR3-GFP (Figure 6B), indicating that DAR3 may function as an inhibitor of DA1. The DA1 peptidase mutant, DA1-pep, showed no effect on reducing wt DA1 peptidase activity on BB-3FLAG cleavage when co-expressed at different levels with DA1 (Figure 6C), indicating that DAR3 had a specific inhibitory effect on DA1 peptidase activity. Co-expression with DAR7 also reduced DA1 peptidase activity, but to a lower extent than DAR3 (Figure 6D).

**Figure 6.**
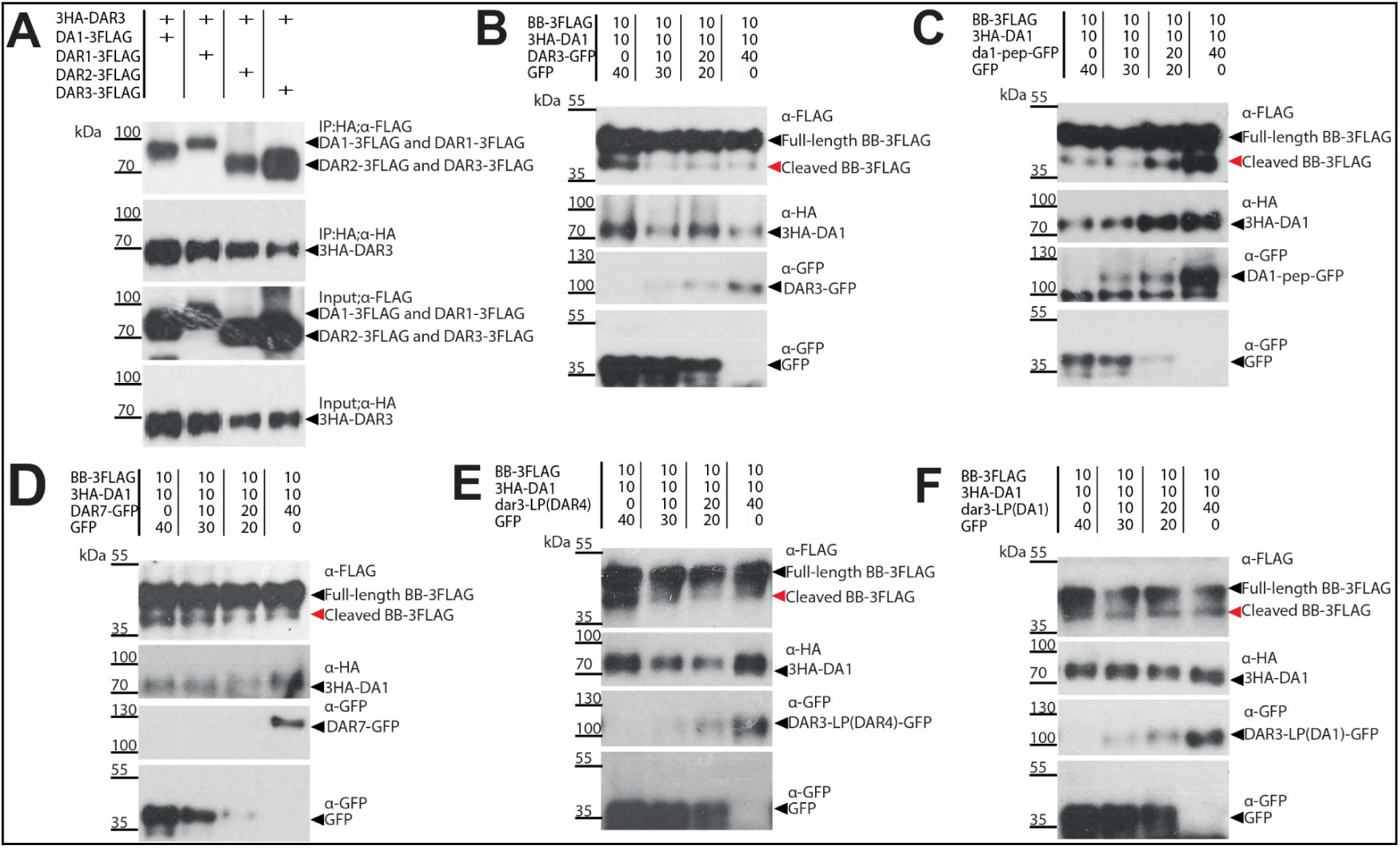
DAR3 and DAR7 family members inhibit DA1 peptidase activity. Immunoblots of Col-0 *da1-ko1dar1-1* transfected mesophyll protoplasts. The numbers above each lane show the fold levels of 10 µg units of transfected DNA *35S::GFP* was used to balance total DNA levels to 60 µg per transfection. The red arrows indicate cleaved BB-3FLAG while the black arrows indicate intact BB-3FLAG. (A) Co-immunoprecipitation of 3HA-DAR3 and DA1-, DAR1- , DAR2 and DAR3-3FLAG. (B) Progressive inhibition of 3HA-DA1 peptidase activity on BB-3FLAG by increasing levels of DAR3-GFP. (C) Increased levels of inactive DA1-peptidase mutant-GFP do not inhibit 3HA-DA1 peptidase activity on BB-3FLAG. (D) DAR7-GFP also progressively inhibits 3HA-DA1 peptidase activity on BB-3-FLAG. (E) Progressive reduction of 3HA-DA1 peptidase activity on BB-3FLAG by DAR3-GFP containing the LP region of DAR4. (F) DAR3-GFP containing the LIM-Peptidase of DA1 does not affect 3HA-DA1 peptidase activity on BB-3FLAG.

As DAR3 is the closest relative of DAR4 (Figure 2A), and because it inhibits DA1 peptidase activity more than DAR7, we assessed the specificity of LIM- Peptidase domains on this inter-family modulation of DA1 peptidase activity. The LIM- Peptidase domains of DA1, DAR3 and DAR4 were swapped and tested for their effects on DA1 peptidase activity. Inhibition of DA1 peptidase activity by DAR3-LP^DAR4^, comprising the LIM- Peptidase domain of DAR4 in DAR3 was compared with DAR3-LP^DA1^ as a control. There was a strong reduction in DA1 peptidase activity by DAR3-LP^DAR4^ compared to DAR3-LP^DA1^ (Figures 6E and 6F). This showed that the LIM- Peptidase domain of DAR4 also inhibited DA1 peptidase activity, consistent with the high degree of sequence conservation between DAR3 and DAR4 LIM- Peptidase domains. These observations establish a link between the inhibition of DA1 peptidase activity by DAR3 and the function of DAR4 as an integrated decoy recognising changes in the level and activities of DAR3 mediated by putative pathogen effectors.

### *DAR3* is involved in resistance to multiple pathogens

As DA1 family peptidase activity promotes pathogen infection and is required for host ploidy increases (Figures 4 and 5), and because DAR3 inhibits DA1 peptidase activities (Figure 6), one plausible mechanism by which pathogens could increase DA1 activity to promote infection involves reducing DAR3-mediated inhibition of DA1. This was tested by expressing *DAR3-GFP* from the *35S* promoter in the susceptible Ws-2 accession and measuring DAR3-GFP protein stability during AcEM2 infection. DAR3-GFP protein levels were rapidly depleted AcEM2- infected but not water-treated plants. DAR3-GFP levels were reduced from 2 dpi, were very low between 4 and 6 dpi, and showed a slight increase between 8 dpi and 10 dpi (Figure 7A). In comparison 3HA-DA1 stability was not affected by AcEm2 infection. AcEm2 growth was tested on *DAR3-GFP* over- expression lines and in the *dar3* CrispR knock- out lines *dar3-2* and *dar3-3* in Ws-2 (Supplementary Figure 9). Over-expression of *DAR3* only delayed AcEM2 growth at 10 dpi, while growth on the *dar3* loss of function lines was similar to wild type (Figure 7B and 7F and Supplementary Table 9). The depletion of DAR3-GFP in over- expression lines by AcEM2 is consistent with the lack of differences in pathogen growth in the *dar3* loss of function lines (Figure 7A).

**Figure 7.**
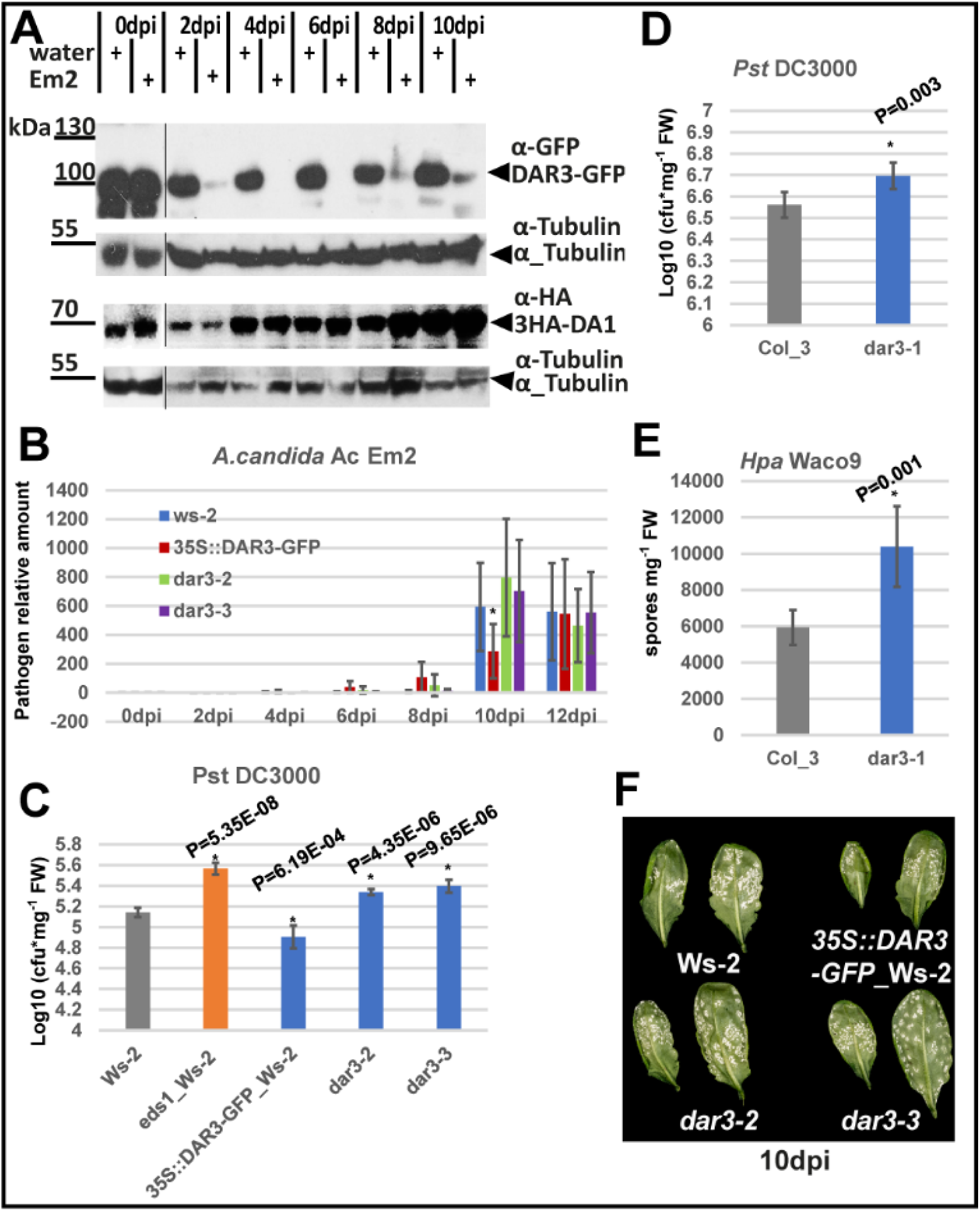
*DAR3* pathogen resistance. (A) The immunoblot shows a time-course of DAR3-GFP and 3HA-DA1 protein stability during *A. candida* Ac EM2 infection of Ws-2 expressing *35S::DAR3-GFP* or *35S::3HA-DA1*. Transgenic seedlings were treated with Em2 spores or water as a control. Tubulin was used as a loading control for plant protein levels. dpi: day post inoculation. The panels on the left showing DAR3-GFP levels at 0 dpi were exposed for a shorter time than the other panel. (B) Time-course of *A. candida* Ac Em2 growth on *Arabidopsis thaliana* Ws-2 and *DAR3* over-expressional and crispr mutant lines. Infection progress was measured using the ratio of pathogen:host DNA levels. dpi days post-infection. (C) *Pst* DC3000 growth on *Arabidopsis thaliana* Ws-2 and *DAR3* over-expressional and crispr mutant lines. cfu: colony-forming units per mg fresh weight of inoculated leaves. The Ws-2 *eds1* mutant was used as a standard susceptible control. (D) (E) The *dar3-1* mutant in Col-3 is more susceptible to *Pst* DC3000 (D) and *Hpa* variety Waco9 (E). Statistical significance was at *P < 0.01 based on a two-tailed Student’s t test. Error bars represent SD of the mean. (F) Image of *A. candida* Ac EM2 growth on leaves of Ws-2 and *DAR3* over-expression and mutant lines at 10 dpi leaves. The white regions are AcEM2.

However, *Pst DC3000* growth was reduced in the Ws-2 *DAR3* over-expression line and increased in the *dar3* loss of function lines (Figure 7C and Supplementary Table 10 ), indicating AcEm2 might carry an effector that targets DAR3. These observations were supported by analyses of *Pst DC3000* and *Hpa* Waco9 growth in a T-DNA *dar3-1* knock-out mutant the background of Col-3 (Supplementary Figure 10 and Table 8). Increased growth of both pathogens was observed when *DAR3* was not functional (Figure 7D and 7E and Supplementary Table 7). These results suggest that *DAR3* reduces the growth of several pathogens and its rapid reduction during early stages of AcEM2 infection facilitates infection.

### DAR3 and DAR4 originate from a common ancestor

These genetic and biochemical analyses establish DAR4 as an integrated domain that function as a decoy for the DAR3 family member that may be a target of *Albugo* effectors. Effector targets might evolve at relatively high rates compared to non-targets [24], therefore DAR3 and DAR4 may be more diverse within the *Brassicaceae* than non-target family members. Phylogenetic analyses of LIM-peptidase diversity in the *Brassicaceae* showed that the DA1 family forms two clades: Clade I comprises three subgroups of DA1, DAR1 and DAR2-like proteins with relatively low diversity; Clade II has three subgroups of DAR3 and DAR4-like, DAR5 and DAR6-like, and DAR7 -like proteins with higher diversity that contain 8 NLR proteins (Figure 8). Five DAR5-like proteins encoded by *B. juncea, B napus, B. oleracea* and *R. sativus* clustered in one branch, suggesting a common origin. In contrast, AtDAR5 was closest to AtDAR6 and clustered with CsDAR6-like proteins, indicating an independent origin from the other five DAR5-like proteins. Two NLR- DAR4-like proteins clustered in one branch.Therefore, at least three LIM-peptidase integrations in Clade II DA1 family proteins, one for DAR4 and two for DAR5, occurred during the evolution and domestication of *Brassicaceae*. Multiple domain integrations and high protein diversity are consistent with the targeting of Clade II proteins by pathogen effectors [24, 29]. In Clade II, the DAR5-like NLR proteins are clustered with DAR6, and two NLR-DAR4-like proteins are clustered with non-NLR-DAR4 and DAR3 proteins. Their common origin and functional conservation is consistent with the evidence that NLR-DAR4 is the R protein detecting effectors targeting DAR3-like inhibitors of DA1 peptidase activity.

**Figure 8.**
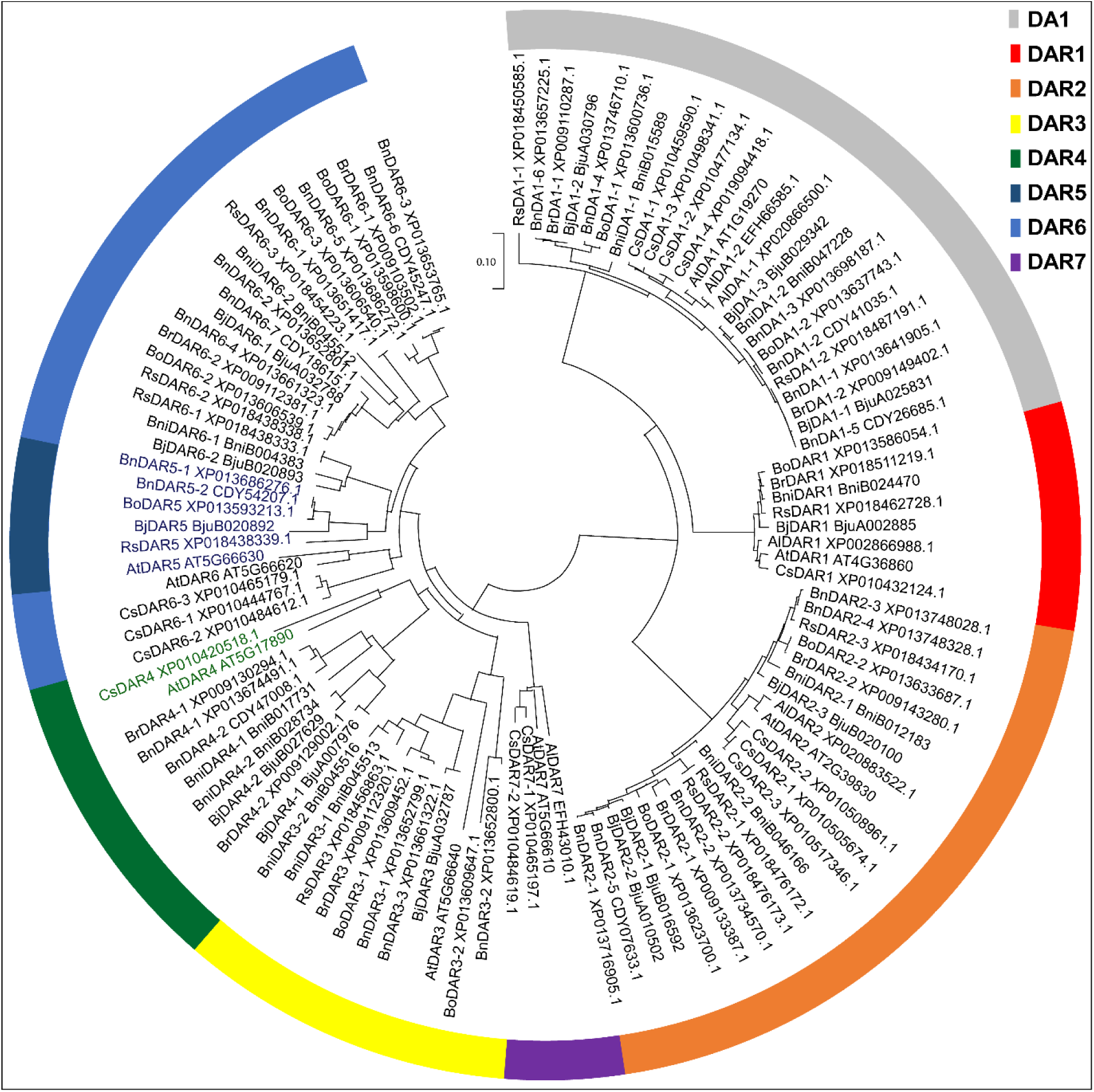
LIM-peptidase domain protein sequence diversity of *Brassicaceae* DA1 family members. The diagram shows a neighbour-joint phylogenetic analysis of DA1 family members identified from sequenced *Brassicaceae* species *Arabidopsis thaliana* (At), *Arabidopsis lyrata* (Al), *Camelina sativa* (Cs, False Flax), *Brassica juncea* (Bj, Brown Mustard), *Brassica napus* (Bn, Oilseed Rape), *Brassica nigra* (Bni, Black Mustard), *Brassica oleracea* (Bo, Chinese Cabbage), *Brassica rapa* (Br, Cabbage) and *Raphanus. sativus* (Rs, Radish). Phylograms were constructed using amino acid sequences of the LIM-peptidase domain aligned with MUSCLE [36]. Branch lengths represent the genetic change between the aligned sequences and circles correspond to bootstrap values >80 following 100 replicates. NLR proteins are indicated by coloured text, green for DAR4 and dark blue for DAR5.

## Discussion

Plant pathogen effectors can promote disease by targeting a wide variety of host proteins. Hosts have evolved immune receptors that detect these interactions, in which effector targets have fused with NLR domains to sense effector activities and trigger defence responses. NLR fusions to LIM-peptidase domains are widespread in several major plant families [9, 10] but their roles in effector sensing and pathogen growth are unknown. Here we show using domain swaps that the LIM-peptidase ID of the NLR CHS3/DAR4 conferring resistance to the oomycete pathogen *Albugo candida* AcEM2 is an integrated decoy for the DA1 family member DAR3. DAR3 interacts with and inhibits the peptidase activities of DA1 family members, and pathogen infection led to rapid reduction of DAR3 levels. This increases DA1 peptidase activity, promoting host tissue endoreduplication and pathogen growth.

### The LIM- Peptidase Integrated Domain of DAR4 functions as a potential decoy for DA1 family members

Mutations disrupting conserved cysteines in DAR4/CHS3 in the 4th Zn-finger of the LIM domain region and a premature stop codon truncating the peptidase active site region promote HR [27, 37] that is dependent on CSA1 [7]. This suggests that the structural and functional integrity of the LIM peptidase ID of DAR4 is required to maintain putative DAR4 sensor and CSA1 executor resistance gene function in a poised state. Swapping the LIM-peptidase domain of DAR4 with those of the other seven family members showed that those from DA1, DAR1, DAR2, DAR3 and DAR7 did not trigger HR, indicating shared features that were able to maintain this poised state and trigger HR in the DAR4/CSA1 context (Figures 1 and 2). It is therefore likely that the LIM-peptidase domain of DAR4 is an ID, the integrity of which is changed by putative effectors that may also influence other DA1 family members except DAR5 and DAR6, which do trigger HR responses when fused to DAR4/CHS3. Reciprocal swaps of small LIM-peptidase domain regions of DA1 and DAR4 show that the full DAR4 LIM- Peptidase domain does not cleave the DA1 substrate BB, but smaller regions of the DAR4 LIM- Peptidase domain in DA1 have peptidase activity (Figure 3), indicating limited functional conservation of the DAR4 LIM-peptidase ID to DA1, DAR1 and DAR2. Therefore DAR3, which is most closely related to DAR4, and/or DAR7 (Figure 2), may be more likely targets for putative effectors that affect LIM-Peptidase integrity. This is supported by the inhibitory function of DAR3-LP^DAR4^.

IDs are thought to arise by recombination and duplication of putative effector target genes with NLR domains [29], indicating that DAR4 most likely arose from DAR3 in the Brassicaceae family [20]presumably selected to combat *Brassica*-specific pathogens such as *Albugo*. Only *Arabidopsis thaliana* and *Camelina sativa* had DAR4-like LIM-peptidase regions fused to an NLR, suggesting specific evolution of resistance to *Albugo candida* in these genera.

The diversity of NLR pairs in *Arabidopsis thaliana* is highest in the *CHS3-DAR4/CSA1* gene pair [24], indicating extensive co-evolution of changes in NLR- LIM peptidase pairs. Analyses of LIM-peptidase ID sequence diversity in *Brassicaceae* showed relatively high levels of diversity not only in DAR4 as expected, but also in DAR3 and DAR7, the effector targets predicted to be sensed by DAR4 (Figure 8). We hypothesise that their high degree of diversity, in contrast to the conservation of DA1, DAR1 and DAR2, is consistent with their being targets of pathogen effectors, and thus subject to diversification and selection in the “arms race” between hosts and pathogens [17, 24].

### DA1 family peptidases facilitate pathogen growth

According to models of intracellular immunity in plants [17], if the DAR4/CHS3 LIM- Peptidase ID functions as a decoy for other members of the DA1 family, then these other family members may facilitate or inhibit pathogen growth and be targets of pathogen effectors. Parasitic and mutualistic biotrophic growth can induce host cell endoreduplication, with ploidy changes accommodating colonization by supporting nutrient production in infected host cells (reviewed by [35]). During organ growth in Arabidopsis the peptidase activities of DA1, DAR1 and DAR2 redundantly limit cell proliferation and promote the transition from mitotic cell divisions to endoreduplication [20, 31]. We linked these DA1 family activities to pathogen growth by showing that the *da1-ko1dar1-1* double mutant, which reduces endoreduplication during normal plant growth, also reduces bacterial and oomycete growth, while over-expression of DA1 promotes bacterial pathogen growth (Figure 5). TCP14 and TCP15, substrates of DA1 peptidase [22], promote mitotic progression. Mutations in TCP14 and TCP15 facilitate powdery mildew infection [35], consistent with pathogen infection increasing DA1 peptidase activity and reducing TCP14 and TCP15 protein levels. Interaction screens also identified several effectors from different pathogens that converge on TCP14 and TCP15 [16, 22], and TCPs form IDs [24], establishing DA1 family functions among this nexus of effector targets. This is consistent with the frequent identification of cell cycle regulatory proteins as interactors with diverse bacterial and fungal effectors [38], emphasising a central role for modulation of host cell cycle functions by pathogens and commensals in plants.

### Reduction of DAR3 levels by pathogen growth promotes DA1 activity

As DAR4 functions as an ID for other members of the DA1 family that promote pathogen growth, it is possible that putative pathogen effectors may modulate the activities of DA1 family members to facilitate pathogen growth, and this is sensed by the DAR4 LIM- Peptidase ID. In uninfected tissues DAR3 likely interacts with DA1 and inhibits its peptidase activity (Figure 6) and in infected tissues DAR3 is rapidly destabilised (Figure 7A). DAR3 does not have Ubiquitin Interaction Motifs (UIMs) like DA1, DAR1 and DAR2, suggesting it may not be an active peptidase, but instead may normally function as an inhibitor of these family members. There is widespread evidence that effectors influence host protein stability [39] and this may account for the destabilisation of DAR3. For example, some effectors have E3 ligase domains that become ubiquitylated after interaction with host E2 enzymes, and some effectors bind to and suppress activities of host E3 ligases. Putative effectors also include F-box adaptor proteins that may re-target host ubiquitylation systems, may function to de-SUMOylate transcription factors and may inhibit proteasome activities [18, 19]. In the sensor model of ID function [17] DAR4 protein levels may be affected by the same effector(s) that reduce DAR3 levels to trigger HR. This was supported by domain-swaps showing the functional equivalence of the LIM-peptidase ID of DAR4 and DAR3.

## Materials and Methods

### Plant growth conditions

*Nicotiana tabacum* plants were grown in long days (16 hours light/ 8 hours dark) at 24°C. *Arabidopsis thaliana* Col-0 and Ws-2 plants were grown in short days (10 hours light/ 14 hours dark) at 20°C. T-DNA mutants and transgenic lines are listed in Supplementary Tables 14 and 15. Ws-2 was crossed with the Col-0 *da1-ko1dar1-1* double mutant and backcross lines were selected in the BC_7_F_2_ population using primers in Supplementary Table 12 to genotype *DA1* and *DAR1* genes.

### Plasmid constructions

Coding sequences of genes were amplified and ligated into pENTR™/D-TOPO® (ThermoFisher, Catalog number K243520) or pDONR207 by in-fusion (Takara, Catalog Number 638909), then transferred into expression plasmids by LR reactions (Invitrogen, Catalog Number 11791-020). The constructions for protein expression in *Escherichia coli* were ligated coding sequence into pETnT by in-fusion [22]. The site-mutation and domain swapped constructs were amplified by PCR and ligated by in-fusion. DAR3 guide RNAs were assembled using GoldenGate into pAGM4723_*ccdB* using FastRed selection. This selection gene was removed after genotyping by backcross with wild type Ws-2. The primers and plant protein expression plasmids used are listed in Supplementary Tables 12 and 13.

### *Agrobacterium*-Mediated Transient Transformation of *N. tabacum* and HR assays

*Agrobacterium* GV3101 strains were grown in LB-medium overnight supplemented with appropriate antibiotics. Cells were harvested and adjusted to OD600 0.5 in resuspension buffer (10 mM MgCl_2_, 10 mM MES pH 5.6), then infiltrated into 4 to 5 weeks old *N. tabacum* leaves[8]. HR reactions were scored after 2 to 4 days according to [28], and 15 panels were scored for each treatment.

### Protein expression in *Arabidopsis* protoplasts

BB cleavage was assessed by co-expressing 35S::BB-3FLAG with 35S::3HA-DA1 and mutant variants in *da1-ko1dar1-1* leaf protoplasts [8, 22]. DAR3 inhibition of DA1 was tested in *da1-ko1dar1-1* leaf protoplast cells with a concentration of 50,000 cells/200 µl and 10 µg of each DNA.

### DA1 domain swap *in vitro* ubiquitination test

DA1 domain swap proteins were expressed in *E. coli* BL21 (DE3) as C-terminal HIS-tagged proteins. For protein expression cells were grown to OD600 0.6 in LB-medium at 37°C, and induced by 0.1mM Isopropyl β- d-1-thiogalactopyranoside (IPTG) for 3 hours at 28°C. Cells were harvested, disrupted by sonication (4 x 5 sec bursts with 20 sec interval) in TGH buffer ( 50 mM HEPES-KOH pH 7.4, 150 mM NaCl, 10% (v/v) glycerol, 1% (v/v) Triton X-100, with EDTA-free protease inhibitor cocktail (Roche Catalog number 11873580001)) and purified by Dynabeads™ His-Tag Isolation and Pulldown (ThermoFisher, Catalog number 10103D). DA1 swapped protein ubiquitylation reactions used E1, E2, E3 with ubiquitin in reaction buffer at 30⁰C for 8 hours *in vitro* according to [22]. Protein ubiquitination was tested by immunoblots and FLAG-HRP antibody (Sigma, Catalog number A8592).

### Pathogen growth tests

*Pseudomonas syringae Pst* DC3000 at an OD600 0.02 were sprayed onto 17 day old *Arabidopsis* seedlings and inoculated plants were harvested 40 hours after inoculation [8, 22]. Leaves were sterilised in 75% ethanol for 15 seconds, washed in water for 1 min and weighted before grinded in 10 mM MgCl_2_. Serial dilutions were plated and grown at 28°C for 2 days and colony-forming units were counted. *Hpa* Noco2 spores were sprayed onto 14 days Arabidopsis seedlings at a concentration of 10,000 spores/ml, then grown at 16⁰C for 7 to 10 days. Spores were washed from plants and concentrations were calculated using a cytometer. *A. candida* AcEM2 spores (10^5^ /ml) were sprayed on 4-5 week- old plants, which were then grown in short day conditions for 10 days after infection. Spore growth was tested by quantitative PCR to measure ratios of host genomic DNA to pathogen DNA ITS (pathogen)/EF1-α (host) [40].

### Protein extraction and analyses

Total protein was extracted in 10 mM Tris HCl pH 7.5, 150 mM NaCl, 0.5 mM EDTA, 10% Glycerol, 0.5% NP-40, EDTA-free Protease Inhibitor Cocktail. DAR3-GFP levels were assessed using immunoblots with GFP antibody (MiltenyiBiotec, Anti-GFP-HRP, Catalog number 130-091-833), 3HA-DA1 by HA-HRP (Sigma, Catalog number 12013819001) and αTubulin levels were used as loading control (α_Tubulin antibody, Monoclonal Anti-α-Tubulin antibody produced in mouse, Sigma, Catalog number T5168; anti-Rabbit IgG, Sigma, Catalog number A0545).

### Protein co-immunoprecipitation

3HA-DAR3 and DA1 family proteins tagged with 3FLAG were expressed in *da1-ko1dar1-1* leaf protoplast cells. Proteins were extracted and purified using Pierce Anti-HA magnetic beads (ThermoFisher Catalog number 88836) for 3 hours. After 4 washes in 10 mM TrisHCl pH 7.5, 150 mM NaCl, 10% Glycerol, 0.5% NP-40, EDTA-free Protease Inhibitor Cocktail, bound proteins were eluted at 95⁰C for 5 minutes in 1 x SDS sample buffer. Protein levels were assessed using immunoblots with HA-HRP (Sigma, Catalog number 12013819001) or FLAG-HRP antibodies (Sigma, Catalog number A8592).

### Ploidy levels

Three 5mm leaf discs from one plant were chopped in Otto buffer I, filtered through a 30 μm filter, diluted with 2 volumes of Otto buffer II and kept at room temperature for 5 min. Ploidy levels were measured using a BD FACSMelody™ Cell Sorter [41]. The ploidy index was calculated as (% 4C nuclei x 1) + (% 8C nuclei x 2) + (% 16C nuclei x 3) + (% 32C nuclei x 4) + (% 64C nuclei x 5).

### Phylogenetic analysis

Proteins were aligned using the MUSCLE algorithm [36] and neighbour-joint trees were created using MEGA-X (https://www.megasoftware.net/) . Protein sequence accessions are listed in Supplementary Tables 1, 2 and 11.

## Author contributions

B.G., V.C., J.D.G.J. and M.W.B. designed the research. B.G., T.P. C.S., F.H.-L., H.D. and N.McK. performed experiments, B.G. analysed data, and M.W.B., BG and J.D.G.J. wrote the manuscript.

## Data Availability

Data generated in this research can be found in the Supplementary Tables.

## Acknowledgments

We thank Dr. Yan Ma for advice on *N. tabacum* infiltration, and Drs. Catherine Gardener, Dae Sung Kim, Xiaokun Liu and Pingtao Ding for advice on pathogen growth tests. Work in the MWB lab was supported by the Biological and Biotechnological Sciences Research Council (BBSRC) Newton Fund to BG and MWB), BBSRC Grant BB/K017225, and Institute Strategic Grant GEN (BB/P013511/1). Work in the JDGJ lab was supported by a Gatsby charitable foundation grant to TSL Norwich, by BBSRC BB/M003809/1 (supported VK) and ERC grant 233376 “Albugon” (supported VK). Work in the VC lab was supported by the University of Bath University Research Studentship Account (URSA), University of Bath start-up fund and Royal Society International Collaboration Awards (ICA\R1\201511).

## Supplementary Files

Supplementary Figure 1. Hypersensitive Response (HR) test of DAR4 auto-immunity mutants.

Supplementary Figure 2. DAR4 and CSA1 form an R protein pair.

Supplementary Figure 3. Protein sequence alignment of three R gene pairs, *DAR4* and *CSA1*, *RRS1* and *RPS4*, and *RRS1B* and *RPS4B*.

Supplementary Figure 4. Expression levels of wild-type and mutant DAR4 proteins in *N. tabacum* leaves.

Supplementary Figure 5. Alignment of LIM-peptidase domain protein regions of DA1 family proteins from *Arabidopsis thaliana* accession Col-0.

Supplementary Figure 6. *In vitro* ubiquitylation of DA1 and domain swapped versions. Supplementary Figure 7. *Albugo candida* Em2 growth in Ws-2 leaves expressing *DA1-GFP* from the 35S promoter.

Supplementary Figure 8. Leaf cell ploidy profiles of *Arabidopsis thaliana* accession Ws-2 lines infected by *Albugo candida* Ac Em2.

Supplementary Figure 9. Designing the *dar3-2* and *dar3-3* CRISPR alleles of *DAR3* in the Ws-2 background.

Supplementary Figure 10. Characterising the *dar3-1* (SAIL_15_D09)T-DNA allele in the Col-3 background.

Supplementary Tables.

