## Supplementary Files for "The integrated LIM-peptidase domain of the CSA1/CHS3 paired immune receptor detects changes in DA1 family peptidase inhibitors to confer *Albugo candida* resistance in Arabidopsis"

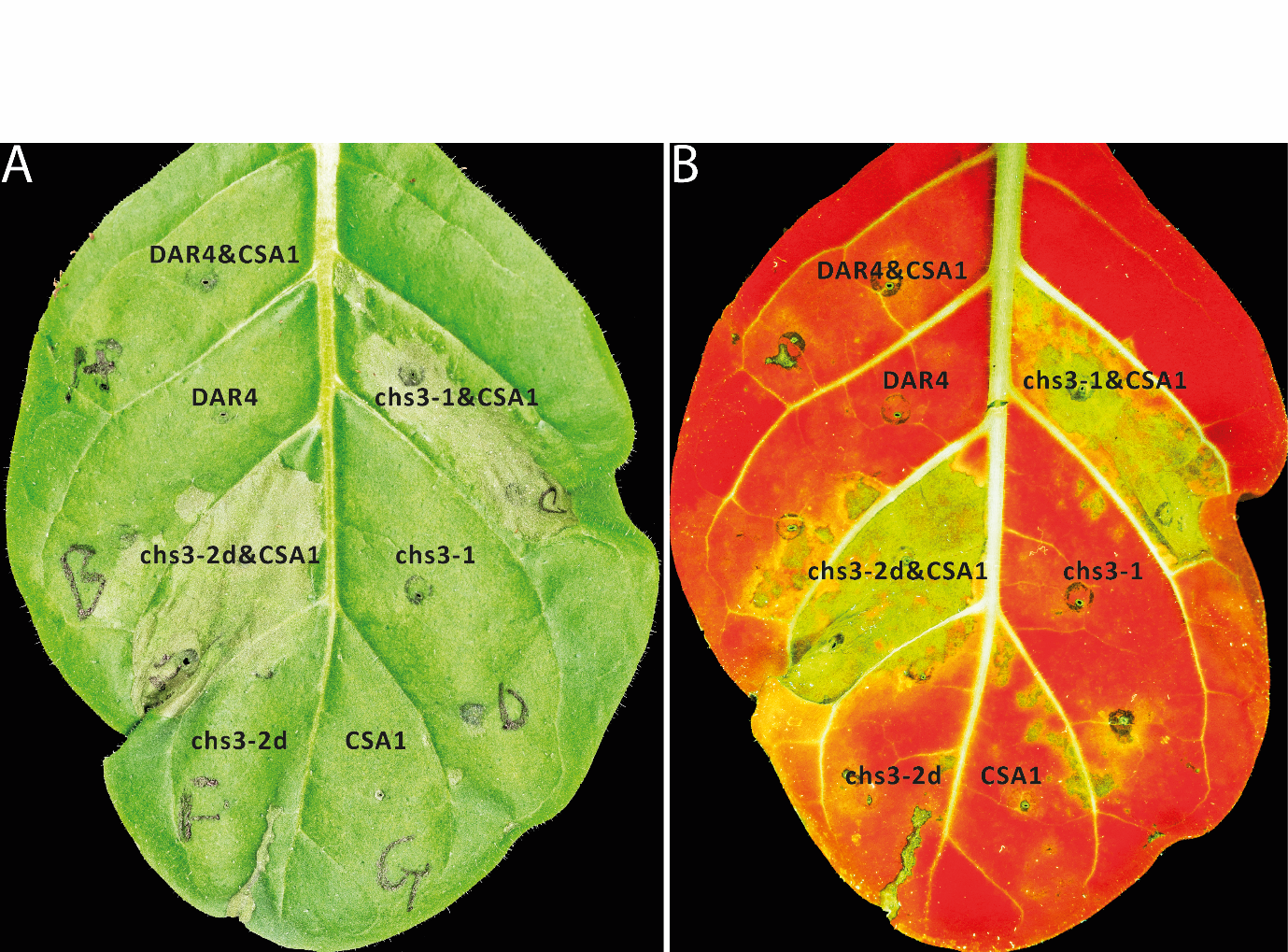

**Supplementary Figure 1. Hypersensitive Responses (HR) of DAR4 auto-immunity mutants in transiently infected N. tabacum leaves.**

(A) Normal light, green leaf indicates no HR and brown leaf indicates HR.

(B) UV light, red region indicates no HR and yellow regions indicate HR.

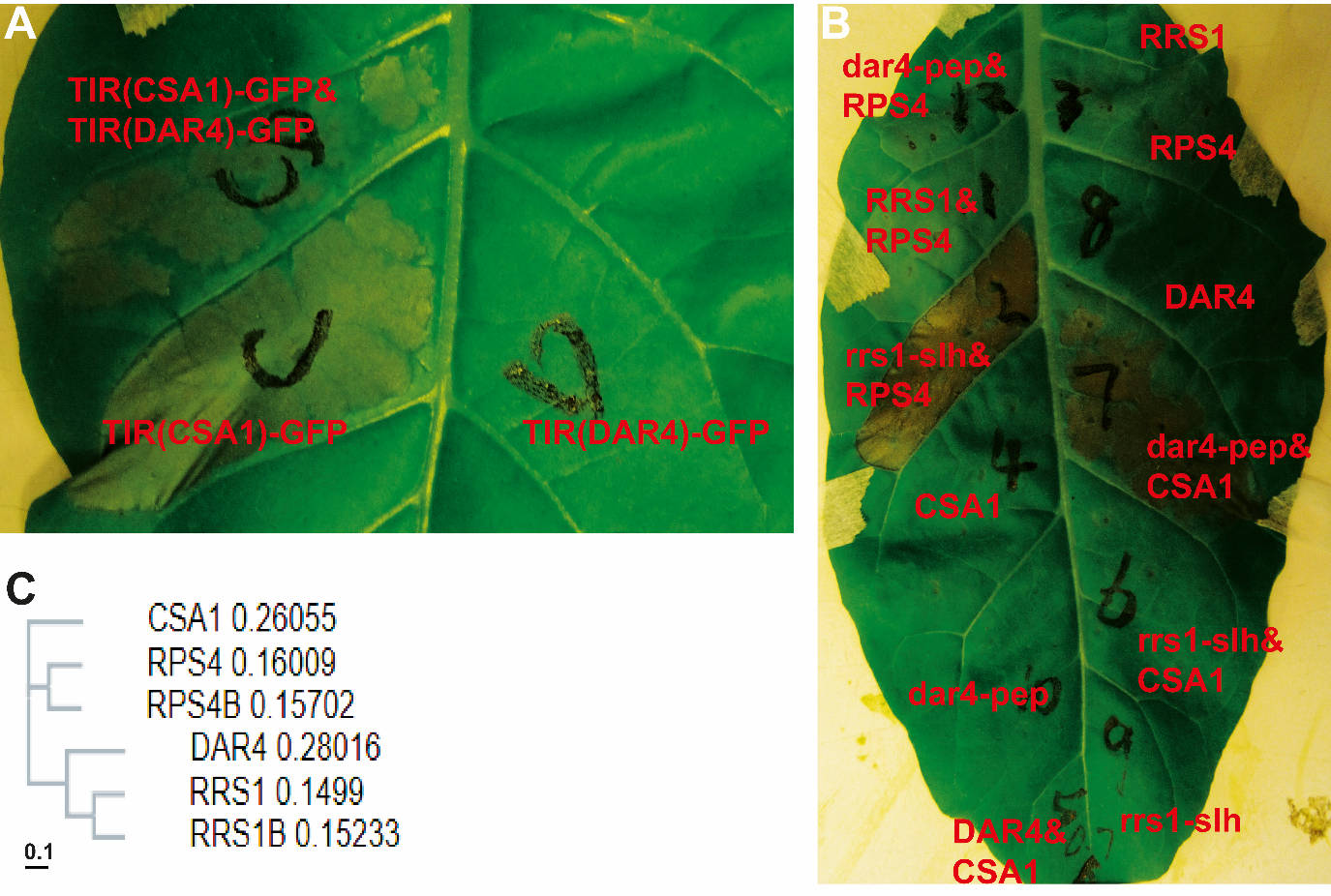

**Supplementary Figure 2. DAR4 and CSA1 form an R protein pair.**

(A) The TIR domain of *CSA1* triggered HR in *N. tabacum* leaves, but the TIR domain of *DAR4* did not. The proteins were tagged by GFP, TIR(DAR4)-GFP and TIR(CSA1)-GFP.

(B) HR test of *DAR4* & *CSA1* and *RRS1* & *RPS4* R gene pairs*. rrs1-slh* is an autoimmune mutant of *RRS1*. *rrs1-slh* & *RPS4* and *dar4-pep* and *CSA1* triggered HR, Mixed R protein pairs, *dar4-pep* & *RPS4* and rrs1-slh&CSA1, did not trigger HR. All controls, including the single proteins DAR4, CSA1, dar4-pep, RPS4, RRS1, *rrs1-slh*, and the native pairs, DAR4 and CSA1,and RRS1 and RPS4, did not trigger HR.

(C) Phylogenetic analysis of the R protein pairs, DAR4 & CSA1, RRS1 & RPS4 and RRS1B & RPS4B. The bar at bottom indicates the distance scale.

```
CSA1       ----MTSSSSWVKTDGETPQDQVFINFRGVELRKNFVSHLEKGLKRKGINAFIDTDEEMG	56
RPS4       -----METSSISTVEDKPPQHQVFINFRGADLRRRFVSHLVTALKLNNINVFIDDYEDRG	55
RPS4B      -------MAASSSSTGLPPQHQVFINFRGEDLRLGFVSHLVEALENDNIKVFIDNYADKG	53
DAR4       MEPPAARVTPSIKADCS---HSVNI-ICEETVLHSLVSHLSAALRREGISVFVDACGLQE	56
RRS1       ------------MTNCEKDEEFVCI-SCVEEVRYSFVSHLSEALRRKGINNVVVDVDIDD	47
RRS1B      ---------------MTESEQIVYI-SCIEEVRYSFVSHLSKALQRKGVNDVFID--SDD	42
                               . * *      :   :****  .*. ..:. ..

CSA1       Q---ELSVLLERIEGSRIALAIFSPRYTESKWCLKELAKMKERTEQ-KELVVIPIFYKVQ	112
RPS4       Q---PLDVLLKRIEESKIVLAIFSGNYTESVWCVRELEKIKDCTDE-GTLVAIPIFYKLE	111
RPS4B      E---PLETLLTKIHDSKIALAIFSGKYTESTWCLRELAMIKDCVEK-GKLVAIPIFYKVD	109
DAR4       TKFFSIKQNQPLTDGARVLVVVISDEVEFYDPWFPKFLKVIQGWQN-NGHVVVPVFYGVD	115
RRS1       L---LFKESQAKIEKAGVSVMVLPGNCDPSEVWLDKFAKVLECQRNNKDQAVVSVLYGDS	104
RRS1B      S---LSNESQSMVERARVSVMILPGNRT---VSLDKLVKVLDCQKN-KDQVVVPVLYGVR	95
                 .      . : : : ::  .       . ::  : :   :    ..: ::*

CSA1       PVTVKELKGDFGDKFRELVKSTDKKTKKEWKEALQYVPFLTGIVLDEKSDEDEVINIIIR	172
RPS4       PSTVRDLKGKFGDRFRSMAKG-DE-RKKKWKEAFNLIPNIMGIIIDKKSVESEKVNEIVK	169
RPS4B      PSTVRGVRGQFGDAFRDLEER-DVIKKKEWKQALKWIPGLIGITVHDKSPESEILNEIVK	168
DAR4       SLTRV-----YG-------------WANSWLEAEKLTSHQSKILSNNVLTDSELVEEIVR	157
RRS1       LLR------------------------DQWLSELDFRGLSRIHQSRKECSDSILVEEIVR	140
RRS1B      SSE------------------------TEWLSALDSKGFSSVHHSRKECSDSQLVKETVR	131
                                       .* .  .           .   :.  ::  ::

CSA1       KVKEILNRRSEGPPSKCSALPPQ-------RHQKRHETFWGIELRIKQLEEKLRFGSDET	225
RPS4       AVKTALTGIPPEGSHNAVVGALG-NSNAGTSSGDKKHETFGNEQRLKDLEEKLDRDKYKG	228
RPS4B      EVKKVLKKVSLEGSQKVVSVDPSQSIDTLSSVGGEKDKTFGIKQRLKELEEKLDLVKYKG	228
DAR4       DVYGKLYPA----------------------------ERVGIYARLLEIE-KLLYKQHRD	188
RRS1       DVYETHFYV----------------------------GRIGIYSKLLEIE-NMVNKQPIG	171
RRS1B      DVYEKLFYM----------------------------ERIGIYSKLLEIE-KMINKQPLD	162
            *                                      *   :: ::* ::   .

CSA1       TRTIGVVGMPGIGKTTLATMLYEKWNDRFLRHVLIRDIHEASEEDGLNYLATKFLQGLL-	284
RPS4       TRIIGVVGMPGIGKTTLLKELYKTWQGKFSRHALIDQIRVKSKHLELDRLPQMLLGE-LS	287
RPS4B      TRVIGVVGMPGIGKTTLVKELYKTWQGKFSRYALIDQIRGKSNNFRLECLPTLLLEKLLP	288
DAR4       IRSIGIWGMPGIGKTTLAKAVFNHMSTDYDASCFIENFDEAFHKEGLHRLLKERIGKILK	248
RRS1       IRCVGIWGMPGIGKTTLAKAVFDQMSSAFDASCFIEDYDKSIHEKGLYCLLEEQLLPG--	229
RRS1B      IRCVGIWGMPGIGKTTLAKAVFDQMSGEFDAHCFIEDYTKAIQEKGVYCLLEEQFLKE--	220
            * :*: ********** . ::.  .  :    :* :     ..  :  *    :

CSA1       KVENANIESVQAAHEAYKDQLLETKVLVILDNVSNKDQVDALLGE------RNWIK---K	335
RPS4       KLNHPHVDNLKDPYSQ----LHERKVLVVLDDVSKREQIDALREI------LDWIKEGKE	337
RPS4B      ELNNPQLDSIEEPYKTHKGLLRERKVLVVLDDVSRREQIYALLGKYDLHSKHEWIK---D	345
DAR4       D-EFDIESSYIMRPTLHRDKLYDKRILVVLDDVRDSLAAESFLKR------LDWFG---S	298
RRS1       ------NDATIMKLSSLRDRLNSKRVLVVLDDVRNALVGESFLEG------FDWLG---P	274
RRS1B      ---NAGASGTVTKLSLLRDRLNNKRVLVVLDDVRSPLVVESFLGG------FDWFG---P	268
                  .            * . ::**:**:*       ::          :*:

CSA1       GSKILITTSDKSLMIQSLVNDTYEVPPLSDKDAIKHFIRYAFDGNEGAAPGPGQGNFPKL	395
RPS4       GSRVVIATSDMSL-TNGLVDDTYMVQNLNHRDSLQLFHYHAFIDDQ---ANPQKKDFMKL	393
RPS4B      GSRIIIATNDISS-LKGLVHDTYVVRQLNHRDGLQLFRYHAFHYDQ---ATPPKVDFMKL	401
DAR4       GSLIIITSVDKQVFAFCQINQIYTVQGLNVHEALQLFSQSVFG-IN-----EPEQNDRKL	352
RRS1       GSLIIITSRDKQVFCLCGINQIYEVQGLNEKEARQLFLLSASIKED-----MGEQNLQEL	329
RRS1B      KSLIIITSKDKSVFRLCRVNQIYEVQGLNEKEALQLFSLCASI-DD-----MAEQNLHEV	322
            * ::*:: * .      :.: * *  *. ::. : *   .    :       : :  ::

CSA1       SKDFVHYTKGNPLALQMLGKELLGKDE-SHWGLKLNALDQHHNSPPGQSICKMLQRVWEG	454
RPS4       SEGFVHYARGHPLALKVLGGELNKKSM-DHWNSKMKKLAQSPS--------PNIVSVFQV	444
RPS4B      SDEFVHYARGHPLALKILGRELYEKNM-KHWETKLIILAQSPT--------TYIGEVVQV	452
DAR4       SMKVIDYVNGNPLALSIYGRELMGKK--SEMETAFFELKHCPP--------LKIQDVLKN	402
RRS1       SVRVINYANGNPLAISVYGRELKGKKKLSEMETAFLKLKRRPP--------FKIVDAFKS	381
RRS1B      SMKVIKYANGHPLALNLYGRELMGKKRPPEMEIAFLKLKECPP--------AIFVDAIKS	374
           *  .:.*..*:***:.: * **  *.   .    :  * .             :  . :

CSA1       SYKALSQKEKDALLDIACFR-SQDENYVASLLDSDGPSN-----ILEDLVNKFMINIYAG	508
RPS4       SYDELTTAQKDAFLDIACFR-SQDKDYVESLLASSDLGSAEAMSAVKSLTDKFLINTCDG	503
RPS4B      SYDELSLAQKDAFLDIACFR-SQDVDYVESLLVSSDPGSAEAI---KALKNKFLIDTCDG	508
DAR4       AYSALSDNEKNIVLDIAFFFKGETVNYVMQLLEESHYFPRLA---IDVLVDKCVLTISEN	459
RRS1       TYDTLSDNEKNIFLDIACFFQGENVNYVIQLLEGCGFFPHVE---IDVLVDKCLVTISEN	438
RRS1B      SYDTLNDREKNIFLDIACFFQGENVDYVMQLLEGCGFFPHVG---IDVLVEKSLVTISEN	431
           :*. *.  :*: .**** *  .:  :** .**              . * :* ::    .

CSA1       KVDMHDTLYMLSKELGREATATDRKGRHRLWHHHTI---------------IAVLDKNKG	553
RPS4       RVEMHDLLYKFSREVDLKASNQDGSRQRRLWLHQHIIK----------GGIINVLQNKMK	553
RPS4B      RVEMHDLLYRFSRELDLKASTQGGSKQRRLWVRQDI---------------INVQQKTMG	553
DAR4       TVQMNNLIQDTCQEIFNGEI----ETCTRMWEPSRIRYLLEYDELEGSGETKAMPKSGLV	515
RRS1       RVWLHKLTQDIGREIINGETV-QIERRRRLWEPWSIKYLLEYNEHKANGEPKTTFKRAQG	497
RRS1B      RVRMHNLIQDVGRQIINRETR-QTKRRSRLWEPCSIKYLLEDKEQNENEEQKTTFERAQV	490
            * ::.      :::         .   *:*    *                   .

CSA1       GSNIRSIFLDLSDITRKWCFYRHAFAMMRDLRYLKIYSTHCPQECESDIKLNFPEG-LLL	612
RPS4       AANVRGIFLDLSEVEDETSLDRDHFINMGNLRYLKFYNSHCPQECKTNNKINIPDK-LKL	612
RPS4B      AANVRGIFLDLSEVKVETSLDREHFKNMRNLRYLKLYNSHCPHECLTNNKINMPDG-LEL	612
DAR4       AEHIESIFLDTSNVK--FDVKHDAFKNMFNLKFLKIYNSCS----KYISGLNFPKG-LDS	568
RRS1       SEEIEGLFLDTSNLR--FDLQPSAFKNMLNLRLLKIYCSNP----EVHPVINFPTGSLHS	551
RRS1B      PEEIEGMFLDTSNLS--FDIKHVAFDNMLNLRLFKIYSSNP----EVHHVNNFLKGSLSS	544
             .:..:*** *::     .    *  * :*: :*:* :            *:    *

CSA1       PLNEVRYLHWLKFPLKEVPQDFNPGNLVDLKLPYSEIERVWEDNKDAPKLKWVNLNHSKK	672
RPS4       PLKEVRCLHWLKFPLETLPNDFNPINLVDLKLPYSEMEQLWEGDKDTPCLRWVDLNHSSK	672
RPS4B      PLKEVRCLHWLKFPLEELPNDFDPINLVDLKLPYSEIERLWDGVKDTPVLKWVDLNHSSK	672
DAR4       LPYELRLLHWENYPLQSLPQDFDFGHLVKLSMPYSQLHKLGTRVKDLVMLKRLILSHSLQ	628
RRS1       LPNELRLLHWENYPLKSLPQNFDPRHLVEINMPYSQLQKLWGGTKNLEMLRTIRLCHSHH	611
RRS1B      LPNVLRLLHWENYPLQFLPQNFDPIHLVEINMPYSQLKKLWGGTKDLEMLKTIRLCHSQQ	604
               :* *** ::**: :*::*:  :**.:.:***::.::    *:   *: : * ** :

CSA1       LNTLAGLGKAQNLQELNLEGCTALKEMHVDMENMKFLVFLNLRGCTSLKSLPEIQLISLK	732
RPS4       LCSLSGLSKAEKLQRLNLEGCTTLKAFPHDMKKMKMLAFLNLKGCTSLESLPEMNLISLK	732
RPS4B      LCSLSGLSKAQNLQRLNLEGC------------------------TSLESLRDVNLTSLK	708
DAR4       LVECDILIYAQNIELIDLQGCTGLQRFPDT-----------------------SQLQNLR	665
RRS1       LVDIDDLLKAENLEVIDLQGCTRLQNFPAA-----------------------GRLLRLR	648
RRS1B      LVDIDDLLKAQNLEVVDLQGCTRLQSFPAT-----------------------GQLLHLR	641
           *     *  *:::: ::*:**                                 .*  *:

CSA1       TLILSGCSKFKTFQVISDKLEALYLDGTAIKELPCDIGR---------------------	771
RPS4       TLTLSGCSTFKEFPLISDNIETLYLDGTAISQLPMNMEK---------------------	771
RPS4B      TLTLSNCSNFKEFPLIPENLKALYLDGTSISQLPDNVGN---------------------	747
DAR4       VVNLSGCTEIKCFSGVPPNIEELHLQGTRIREIPIFNATHPPKVKLDRKKLWNLL-----	720
RRS1       VVNLSGCIKIKSVLEIPPNIEKLHLQGTGILALPVSTVKP------NHRELVNFLTEIPG	702
RRS1B      VVNLSGCTEIKSFPEIPPNIETLNLQGTGIIELPLSIVKP------NYRELLNLLAEIPG	695
           .: **.*  :* .  :  ::: * *:** *  :*

CSA1       ---------------------------LQRLVMLNMKGCKKLKRLPDSLGQLKALEELIL	804
RPS4       ---------------------------LQRLVVLNMKDCKMLEEIPGRVGELKALQELIL	804
RPS4B      ---------------------------LKRLVLLNMKDCKVLETIPTCVSELKTLQKLVL	780
DAR4       -ENFSDVEHIDLECVTNLATVTSNNHVMGKLVCLNMKYCSNLRGLPDMVS----------	769
RRS1       LSE-------ELERLTSLLESNSSCQDLGKLICLELKDCSCLQSLPNMANL--DLNVLDL	753
RRS1B      LSGVSNLEQSDLKPLTSLMKISTSYQNPGKLSCLELNDCSRLRSLPNMVN----------	745
                                        :*  *::: *. *. :*   .

CSA1       SGCSKLNEFPETWGNMSRLEILLLDETAIKDMPKILSVRRLCLNKNEKISRLPDLLNKFS	864
RPS4       SDCLNLKIFPEI--DISFLNILLLDGTAIEVMPQLPSVQYLCLSRNAKISCLPVGISQLS	862
RPS4B      SGCSKLKEFPEI--NKSSLKILLLDGTSIKTMPQLPSVQYLCLSRNDHLIYLPAGINQVS	838
DAR4       ----------------------------------------------------------LE	771
RRS1       SGCSSLNSI---QGFPRFLKQLYLGGTAIREVPQLPQSLEILNAHGSCLRSLPN-MANLE	809
RRS1B      ----------------------------------------------------------LE	747
                                                                     ..

CSA1       QLQWLHLKYCK--------------------NLTHVPQLPPNLQYLNVHGCSSLKTVAKP	904
RPS4       QLKWLDLKYCT--------------------SLTSVPEFPPNLQCLDAHGCSSLKTVSKP	902
RPS4B      QLTRLDLKYCT--------------------KLTYVPELPPTLQYLDAHGCSSLKNVAKP	878
DAR4       SLKVLYLSGCSELEKIMGFPRNLKKLYVGGTAIRELPQLPNSLEFLNAHGCKHLKSINL-	830
RRS1       FLKVLDLSGCSELETIQGFPRNLKELYFAGTTLREVPQLPLSLEVLNAHGS---------	860
RRS1B      LLKALDLSGCSELETIQGFPRNLKELYLVGTAVRQVPQLPQSLEFFNAHGCVSLKSIRL-	806
            *  * *. *.                     :  :*::* .*: ::.**.

CSA1       LVCSIPMKHVNSSFIFTNCNELEQAAKEEIVVYAE---------RKCHLLASALK-----	950
RPS4       LARIMPTEQNHSTFIFTNCENLEQAAKEEITSYAQ---------RKCQLLSYARK-----	948
RPS4B      LARIMSTVQNHYTFNFTNCGNLEQAAKEEITSYAQ---------RKCQLLSDARK-----	924
DAR4       -----DFEQLPRHFIFSNCYRFSSQVIAEFVEKGLVASLA--------------------	865
RRS1       -----DSEKLPMHYKFNNFFDLSQQVVNDFLLKTLTYV--KHIPRG-----Y--------	900
RRS1B      -----DFKKLPVHYTFSNCFDLSPQVVNDFLVQAMANVIAKHIPRERHVTGFSQKTVQRS	861
                   :    : *.*   :.  .  ::

CSA1       --RCDESCVPEILFCTSFPGCEMPSWFSHDAIGSMVEFELPPHWNHNRLSGIALCVVVSF	1008
RPS4       --RYNGGLVSESLFSTCFPGCEVPSWFCHETVGSELEVKLLPHWHDKKLAGIALCAVVSC	1006
RPS4B      --HYNEG--SEALFSTCFPGCEVPSWFGHEAVGSLLQRKLLPHWHDKRLSGIALCAVVSF	980
DAR4       -RAKQEELIKAPEVIICIPMDTRQRSSFRLQAGRNAMTD-LVPWMQKPISGFSMSVVVSF	923
RRS1       ---TQELINKAPTFSFSAPSHTNQNATFDLQSGSSVMTR-LNHSWRNTLVGFGMLVEVAF	956
RRS1B      SRDSQQELNKTLAFSFCAPSHANQNSKLDLQPGSSSMTR-LDPSWRNTLVGFAMLVQVAF	920
               :        .  . *             *             : : *:.: . *:

CSA1       KNCKSHANLI-VKFSCEQNNGEGSSSSI-TWKVGSLIEQDNQEETVESDHVFIGYTNCLD	1066
RPS4       LDPQDQVSRLSVTCTFKVKDEDKSWVAY-TCPVGSWTRHGGGKDKIELDHVFIGYTSCPH	1065
RPS4B      PDSQDQLSCFSVTCTFKIKAEDKSWVPF-TCPVGIWTREGNKKDRIESDHVFIAYISSPH	1039
DAR4       QDDYHNDVGLRIRCVGTWKTWNNQPDRIVERFFQCWAP-TEA-PKVVADHIFVLYDTKMH	981
RRS1       PEDYCDATDVGISCVCRWSNKEGRSCR-IERKFHCWAP-WQVVPKVRKDHTFVFSDVNMR	1014
RRS1B      SEGYCDDTDFGISCVCKWKNKEGHSHR-REINLHCWAL-G---KAVERDHTFVFFDVNMR	975
            :   .   . :      .  :          .            :  ** *:

CSA1       FIKLVKGQGGPKCAPTKASLEFSVRTGT--GGEATLEVLKSGFSFVFEPEENRVPSPRND	1124
RPS4       TIKCHEEGNSDECNPTEASLKFTVTGGT--SENGKYKVLKCGLSLVYAKDKDKNSALETK	1123
RPS4B      SIRCLEEKNSDKCNFSEASLEFTVTSDT--SGIGVFKVLKCGLSLVYENDKNKNSSLEAK	1097
DAR4       PS--DSEENHISMWAHEVKFEFHTVSGENNPLGASCKVTECGVEVITAATGDTSVSGIIR	1039
RRS1       PS--TGEGNDPDIWAGLVVFEFFPINQQTKCLNDRFTVRRCGVRVINVATGNTSLENIAL	1072
RRS1B      PD--TDEGNDPDIWADLVVFEFFPVNKQRKPLNDSCTVTRCGVRLITAVNCNTSIENISP	1033
                   .  .     . ::*               * ..*. .:     :

CSA1       DVK-------GKVKIN--------KTP-------------------------------SA	1138
RPS4       Y----------DMLIG--------------------------------------------	1129
RPS4B      Y----------DVPVE--------------------------------------------	1103
DAR4       ESETITIIEKEDTIIDEEDTPLLSRKPEETNRSRSSSELQKLSSTSSKVRSKGNVFWKWL	1099
RRS1       VL---------------------SLDPVEVSGY---E-VLRVS--YDDLQEMDKVLFLYI	1105
RRS1B      VL---------------------SLDPMEVSGNEDEE-VLRVR--YAGLQEIYKALFLYI	1069


CSA1       NGCFKDQAKGN----ESPKGQWQTYIENS--------------------STNIPSEAHSS	1174
RPS4       -KSFQETSEGVDGRVKKTKGKYVMPVEKNFQETTEGVDGRVNKKKKTRMDNGRPKK---K	1185
RPS4B      -VSFQEPEHGIMEEER--------------------------YINKRRSDDRRPKK---K	1133
DAR4       -GCFPLQPKNLRSRSRRTT-ALEEALEEALKE-----------RE--KLEDTRELQIALI	1144
RRS1       ASLFNDEDVDFVAPLI-AG--IDLDVSSGLKVL--------ADVSLISVSSNGEIVMHSL	1154
RRS1B      AGLFNDEDVGLVAPLI-AN-IIDMDVSYGLKVL--------AYRSLIRVSSNGEIVMHYL	1119
              *     .                                       .

CSA1       QKTG---------FNGF------NGMY---SVCVL--YEMYSH-----------------	1197
RPS4       QRSG---------RDDN------QTRMQVELQ-EGNINSVIMHTVK---NF---------	1217
RPS4B      RKTK---------RDDI------MIISTVTQTCVPSVNARIEDKVT---G----------	1165
DAR4       ESKKIKKIKQADERDQIKHADEREQRKHSKDHEEEEIESNEKEERRHSKDYVIEELVLKG	1204
RRS1       QR---------QMGKEILHGQSMLLSD-----CE----SSMTEN-----L----SDVPKK	1187
RRS1B      LR---------QMGKEILHTESKKTDK-----LVDNIQSSMIATK----E----IEITRS	1157
                         .

CSA1       ------------------------------------------------------------	1197
RPS4       ------------------------------------------------------------	1217
RPS4B      ------------------------------------------------------------	1165
DAR4       KGKRKQLDDDKADEKEQIKHSKDHVEEEVNPPLSKCKDCKSAIEDGISINAYGSVWHPQC	1264
RRS1       KK----------------KHSESRVKKV--------V-----S---IPAIDEGDLWTWRK	1215
RRS1B      KS----------------RRKNNKEKRV------VCV-----V---DRGSRSSDLWVWRK	1187


CSA1       ------------------------------------------------------------	1197
RPS4       ------------------------------------------------------------	1217
RPS4B      ------------------------------------------------------------	1165
DAR4       FCCLRCREPIAMNEISDLRGM------YHKPCYKELR------HPNCYVCEKKIPRTAEG	1312
RRS1       YGQ----KDILG--SRFPRGYYRCAYKFTHGCKATKQVQRSETDSNML--------AITY	1261
RRS1B      YGQ----KPIKS--SPYPRSYYRCAS--SKGCFARKQVERSRTDPNVS--------VITY	1231


CSA1       ------------------------------------------------------------	1197
RPS4       ------------------------------------------------------------	1217
RPS4B      ------------------------------------------------------------	1165
DAR4       LKYHEHPFWMETYCPS-HDGDGTPKCCSCERLEHCGTQYVMLADFRWLCRECMDSAIMDS	1371
RRS1       LSEHNHPRPTKRKALADSTRSTSSSIC---------------------------------	1288
RRS1B      ISEHNHPFPTLRNTLAGSTRSSSSKCSDVTTSASST---VS-QD-----KEGPDKSHLPS	1282


CSA1       ------------------------------------------------------------	1197
RPS4       ------------------------------------------------------------	1217
RPS4B      ------------------------------------------------------------	1165
DAR4       DECQ-PLH-----FEIREFFEGLHMKIEEEFPVYL---VEKNALNKAEKEEKIDKQGDQC	1422
RRS1       ------------------------------------------------------------	1288
RRS1B      SPASPPYAAMVVKEEDMEQWDNMEFDVDVEEDTFIPELFPEDTFADM---DKLEENSQTM	1339


CSA1       ------------------------------------------------------------	1197
RPS4       ------------------------------------------------------------	1217
RPS4B      ------------------------------------------------------------	1165
DAR4       ---------LMVVRGICLSEEQIVTSVSQGVRRMLNKQILDTVTESQRVVRKCEVTAILI	1473
RRS1       ------------------------------------------------------------	1288
RRS1B      FLSRRSSGGNMEAQGKNSSDDREVN---------LPSKILNR------------------	1372


CSA1       ------------------------------------------------------------	1197
RPS4       ------------------------------------------------------------	1217
RPS4B      ------------------------------------------------------------	1165
DAR4       LYGLPRLLTGYILAHEMMHAYLRLNGYRNLNMVLEEGLCQVLGYMWLECQTYVFDTATIA	1533
RRS1       ------------------------------------------------------------	1288
RRS1B      ------------------------------------------------------------	1372


CSA1       ------------------------------------------------------------	1197
RPS4       ------------------------------------------------------------	1217
RPS4B      ------------------------------------------------------------	1165
DAR4       SSSSSSRTPLSTTTSKKVDPSDFEKRLVNFCKHQIETDESPFFGDGFRKVNKMMASNNHS	1593
RRS1       ------------------------------------------------------------	1288
RRS1B      ------------------------------------------------------------	1372


CSA1       --------------------	1197
RPS4       --------------------	1217
RPS4B      --------------------	1165
DAR4       LKDTLKEIISISKTPQYSKL	1613
RRS1       --------------------	1288
RRS1B      --------------------	1372
```

**Supplementary Figure 3. Protein sequence alignment of three R gene pairs, *DAR4* and *CSA1*, *RRS1* and *RPS4*, and *RRS1B* and *RPS4B*.**

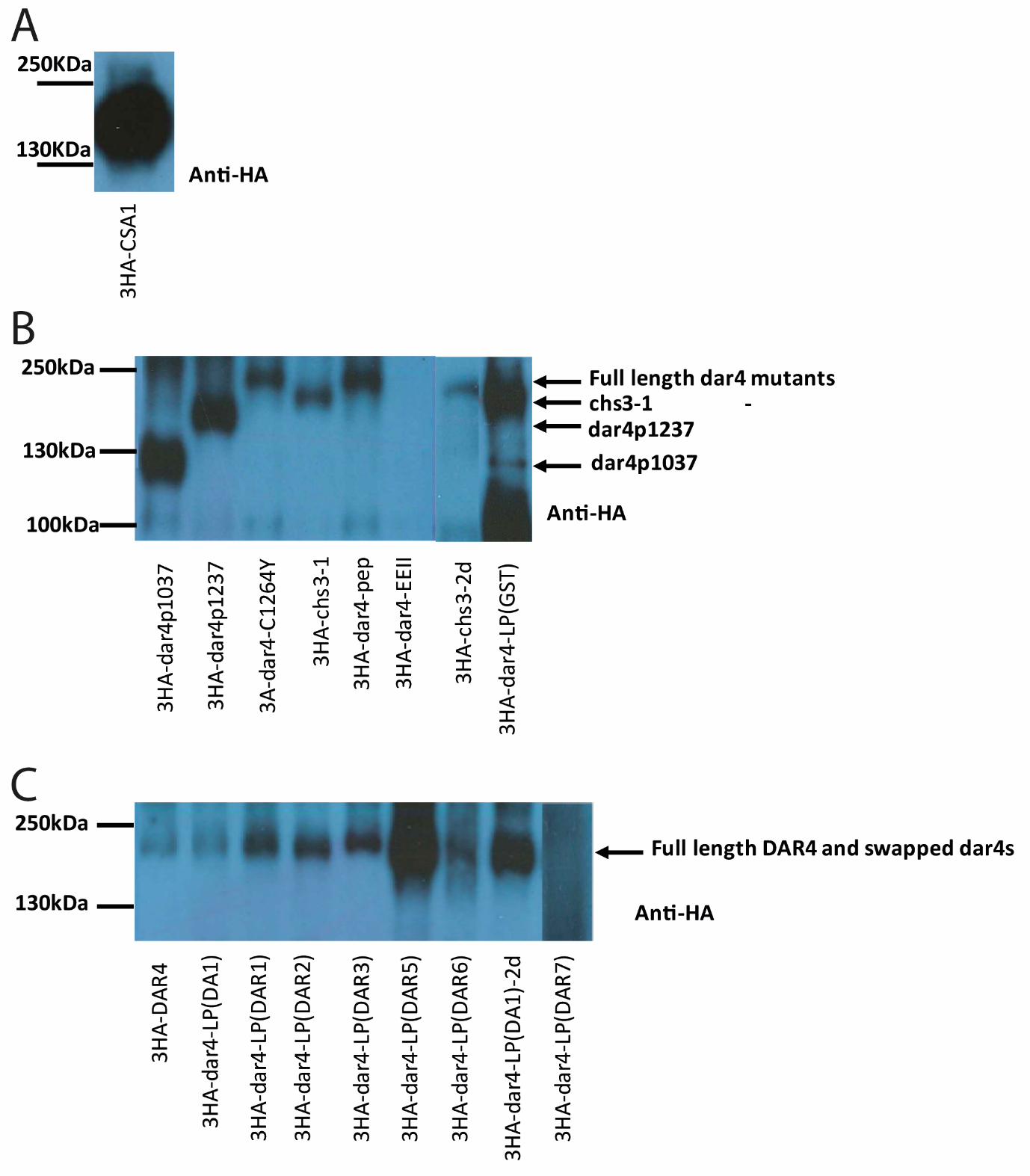

**Supplementary Figure 4. Expression levels of wild-type and mutant DAR4 proteins in *N. tabacum* leaves.**

Immunoblots of crude extracts of *Agrobacterium*- inoculated leaves. 3HA-DAR4 proteins were detected using anti-HA HRP antibodies.

(A) 3HA-CSA1 expression levels

(B) Expression of 3HA-DAR4 mutant proteins. 3HA-dar4-EEII was not detected in these conditions.

(C) Expression levels of 3HA-DAR4 LIM-peptidase domain swapped proteins. 3HA-dar4-LP(DAR7) was not detected in these conditions.

```
DAR2      ---RICGG-CNSDIGSGNYLGCMGTFFHPECFRCHSCGYAITEHEFSL---SGTKPYHKL	212
DA1       ---RICAG-CNMEIGHGRFLNCLNSLWHPECFRCYGCSQPISEYEFST---SGNYPFHKA	222
DAR1      ---RICVG-CQAEIGHGRFLSCMGGVWHPECFCCNACDKPIIDYEFSM---SGNRPYHKL	240
DAR3      SKDVVEEDVNPPPSIDGKSEIGDGTSVNPRCLCCFHCHRPFVMHEIL-----KKGKFHID	120
DAR4      ---SKCKD-CKSAIEDGISINAYGSVWHPQCFCCLRCREPIAMNEIS----DLRGMYHKP	1289
DAR7      ---SICDG-CKSAIEYGRSVHALGVNWHPECFCCRYCDKPIAMHEFS----NTKGRCHIT	250
DAR6      ---SLCGG-CNFAVEHGGSVNILGVLWHPGCFCCRACHKPIAIHDIENHVSNSRGKFHKS	336
DAR5      ---SMCGG-CNSAVKHEESVNILGVLWHPGCFCCRSCDKPIAIHELENHVSNSRGKFHKS	400
                 .               .   :* *: *  *   :   ::           *

DAR2      CFKELT-HPKCEVCHHFIPTNDAGLIEYRCHPFWNQKYCPSHEYDKTARCCSCERLESWD	271
DA1       CYRERY-HPKCDVCSHFIPTNHAGLIEYRAHPFWVQKYCPSHEHDATPRCCSCERMEPRN	281
DAR1      CYKEQH-HPKCDVCHNFIPTNPAGLIEYRAHPFWMQKYCPSHERDGTPRCCSCERMEPKD	299
DAR3      CYKEYYRNRNCYVCQQKIPVNAEGIRKFSEHPFWKEKYCPIHDEDGTAKCCSCERLEPRG	180
DAR4      CYKELR-HPNCYVCEKKIPRTAEG-LKYHEHPFWMETYCPSHDGDGTPKCCSCERLEHCG	1347
DAR7      CYER-S-HPNCHVCKKKFP----G-RKYKEHPFWKEKYCPFHEVDGTPKCCSCERLEPWG	303
DAR6      CYER-Y----CYVCKEK-K----M-KTYNNHPFWEERYCPVHEADGTPKCCSCERLEPRE	385
DAR5      CYER-Y----CYVCKEK-K----M-KTYNIHPFWEERYCPVHEADGTPKCCSCERLEPRG	449
          *:..      * ** .           :  **** : *** *: * * :******:*

DAR2      VRYYTLEDGRSLCLECMETAITDTGECQPLYHAIRDYYEGMYMKLDQQIPMLLVQREALN	331
DA1       TRYVELNDGRKLCLECLDSAVMDTMQCQPLYLQIQNFYEGLNMKVEQEVPLLLVERQALN	341
DAR1      TKYLILDDGRKLCLECLDSAIMDTHECQPLYLEIREFYEGLHMKVEQQIPMLLVERSALN	359
DAR3      TNYVMLGDFRWLCIECMGSAVMDTNEVQPLHFEIREFFEGLFLKVDKEFALLLVEKQALN	240
DAR4      TQYVMLADFRWLCRECMDSAIMDSDECQPLHFEIREFFEGLHMKIEEEFPVYLVEKNALN	1407
DAR7      TKYVMLADNRWLCVKCMECAVMDTYECQPLHFEIREFFGSLNMKVEKEFPLLLVEKEALK	363
DAR6      SNYVMLADGRWLCLECMNSAVMDSDECQPLHFDMRDFFEGLNMKIEKEFPFLLVEKQALN	445
DAR5      TKYGKLSDGRWLCLECGKS-AMDSDECQPLYFDMRDFFESLNMKIEKEFPLILVRKELLN	508
           .*  * * * ** :*      *: : ***:  ::::: .: :*::::. . **.:. *:

DAR2      DAIVGEKNGYHH---MPETRGLCLSEEQTVTSVLRRPRLGAHR-LVGMRTQPQRLTRKCE	387
DA1       EAREGEKNGHYH---MPETRGLCLSEEQTVSTVRKRSKHGTGK-WAGNITEPYKLTRQCE	397
DAR1      EAMEGEKHGHHH---LPETRGLCLSEEQTVTTVLRRPRIGAGYKLIDMITEPCRLIRRCE	416
DAR3      KAEEEEKIDYHR---AAVTRGLCMSEEQIVPSIIKGPRMGPDNQLITDIVTESQRVSGFE	297
DAR4      KAEKEEKIDKQGDQCLMVVRGICLSEEQIVTSVSQGVRRMLNKQILDTVTESQRVVRKCE	1467
DAR7      KAEAQEKIDNQH---GVVTRGICLSEGQIVNSVFKKPTMGPNGELVSLGTEPQKVVGGCE	420
DAR6      KAEKEEKIDYQY---EVVTRGICLSEEQIVDSVSQRPVRGPNNKLVGMATESQKVTRECE	502
DAR5      K--KEEKIDNHY---EVLIRAYCMSEQKIMTYVSEEPRTGQNKQLIDMDTEPQGVVHECK	563
          .    ** .          *. *:** : :  : .              .         :

DAR2      VTAILVLYGLPRLLTGAILAHELMHGWLRLNGFRNLNPEVEEGICQVLSYMWLESEVLSD	447
DA1       VTAILILFGLPRLLTGSILAHEMMHAWMRLKGFRTLSQDVEEGICQVMAHKWLDAELAAG	457
DAR1      VTAILILYGLPRLLTGSILAHEMMHAWLRLNGYPNLRPEVEEGICQVLAHMWLESETYAG	476
DAR3      VTGILIIYGLPRLLTGYILAHEMMHAWLRLNGYKNLKLELEEGLCQALGLRWLESQTFAS	357
DAR4      VTAILILYGLPRLLTGYILAHEMMHAYLRLNGYRNLNMVLEEGLCQVLGYMWLECQTYVF	1527
DAR7      VTAILILYGLPRLLTGYILAHEMMHAWLRLNGYRNLKLELEEGICQVLGHMWLESQTYSS	480
DAR6      VTAILILYGLPRLLTGYILAHEMMHAYLRLNGHRNLNNILEEGICQVLGHLWLDSQTYAT	562
DAR5      VTAILILYGLPRLLTGYILAHEMMHAWLRLNGHMNLNNILEEGICQVLGHLWLESQTYAT	623
          **.**:::******** *****:**.::**:*. .*   :***:**.:.  **:.:

DAR2      PSTRNLPST------SSVATSSSSSFSNKKGGKSNVEKKLGEFFKHQIAHDASPAYGGGF	501
DA1       STNSNAASS------SS------SSQGLKKGPRSQYERKLGEFFKHQIESDASPVYGDGF	505
DAR1      STLVDIASS------SSSA---VVSASSKKGERSDFEKKLGEFFKHQIESDSSSAYGDGF	527
DAR3      TDAAAAAAVASSSSFSSSTAPPAAITSKKSDDWSIFEKKLVEFCMNQIKEDDSPVYGLGF	417
DAR4      DTATIA---S---SSSSSRTPLSTT-TSKKVDPSDFEKRLVNFCKHQIETDESPFFGDGF	1580
DAR7      SAAAS--------SASSSSRTPAAN-ASKKGAQSDYEKKLVEFCKDQIETDDSPVYGVGF	531
DAR6      ADATADASSS---ASSSSRTPPAAS-ASKKGEWSDFDKKLVEFCKNQIETDDSPVYGLGF	618
DAR5      ADTTADAASA---SSSSSRTPPAAS-ASKKGEWSDFDKKLVEFCKNQIETDESPVYGLGF	679
                         **           *.   *  :::* :*  .**  * *  :* **

DAR2      RAANAAACK--YGLRRTLDHIRLTGTFPL----	528
DA1       RAGRLAVHK--YGLRKTLEHIQMTGRFPV----	532
DAR1      RQGNQAVLK--HGLRRTLDHIRLTGTFP-----	553
DAR3      KQVYEMMVSNNYNIKDTLKDIVSASNATPDSTV	450
DAR4      RKVNKMMASNNHSLKDTLKEIISISKTPQYSKL	1613
DAR7      RKVNQMVSD--SSLHKILKSIQHWTKPDSNL--	560
DAR6      RTVNEMVTN--SSLQETLKEILRQR--------	641
DAR5      RTVNEMVTN--SSLQETLKEILRRR--------	702
          :       .   .::  *. *
```

**Supplementary Figure 5. Alignment of LIM-peptidase domain protein regions of DA1 family proteins from *Arabidopsis thaliana* accession Col-0.**

Green highlights conserved amino acids of the 4 zinc-finger stems in the LIM and LIM-like domains. Red highlights the active site region AHEMMHA---EE in the peptidase domain.

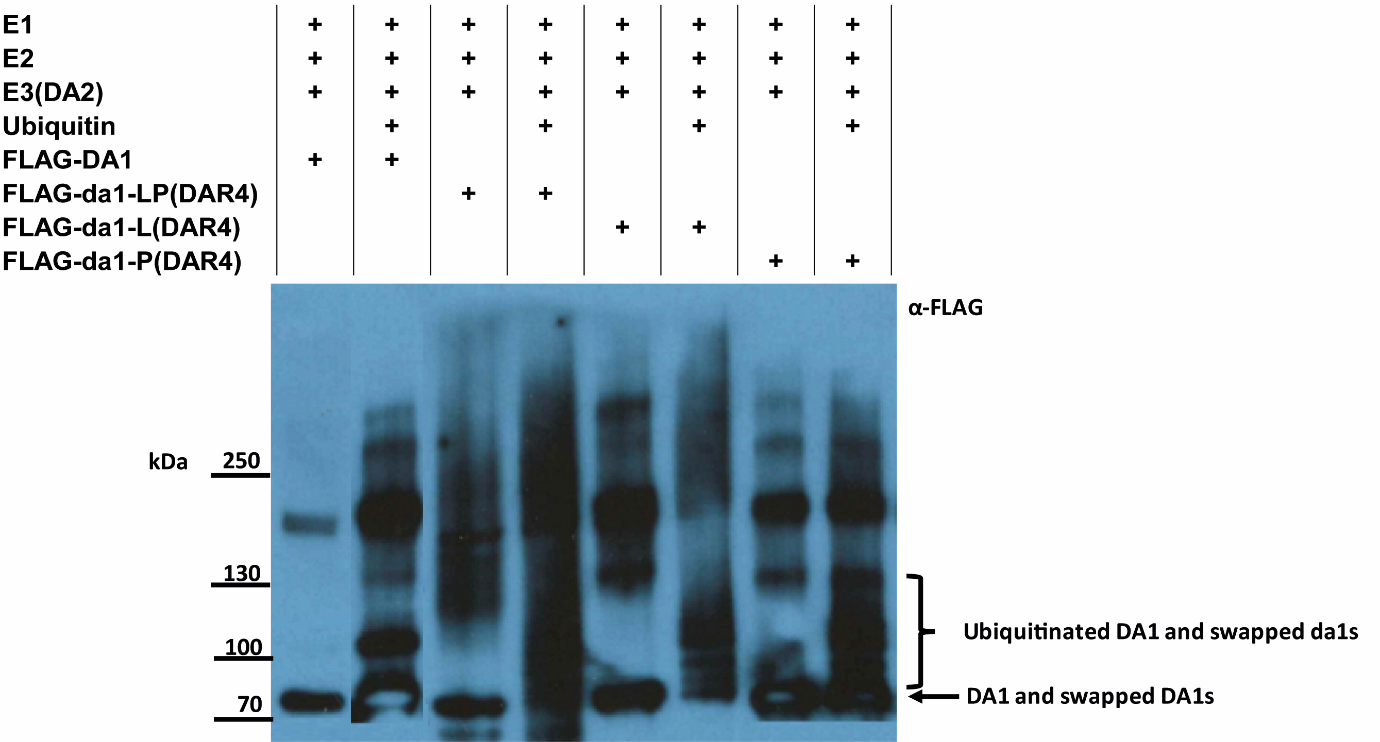

**Supplementary Figure 6. *In vitro* ubiquitylation of DA1 and domain swapped versions.**

*In vitro* reactions contained E1 (human UBE1), E2 (human), E3 (DA2-HIS, expressed in *E. coli* and purified using affinity resin), and ubiquitin (human recombinant). DA1 and domain swapped version proteins were tagged with 3- FLAG at N terminal, expressed in *E. coli* and purified using affinity resin. All human proteins were obtained from Boston Biochemical.

Ubiquitylated 3FLAG-DA1 proteins are identified.

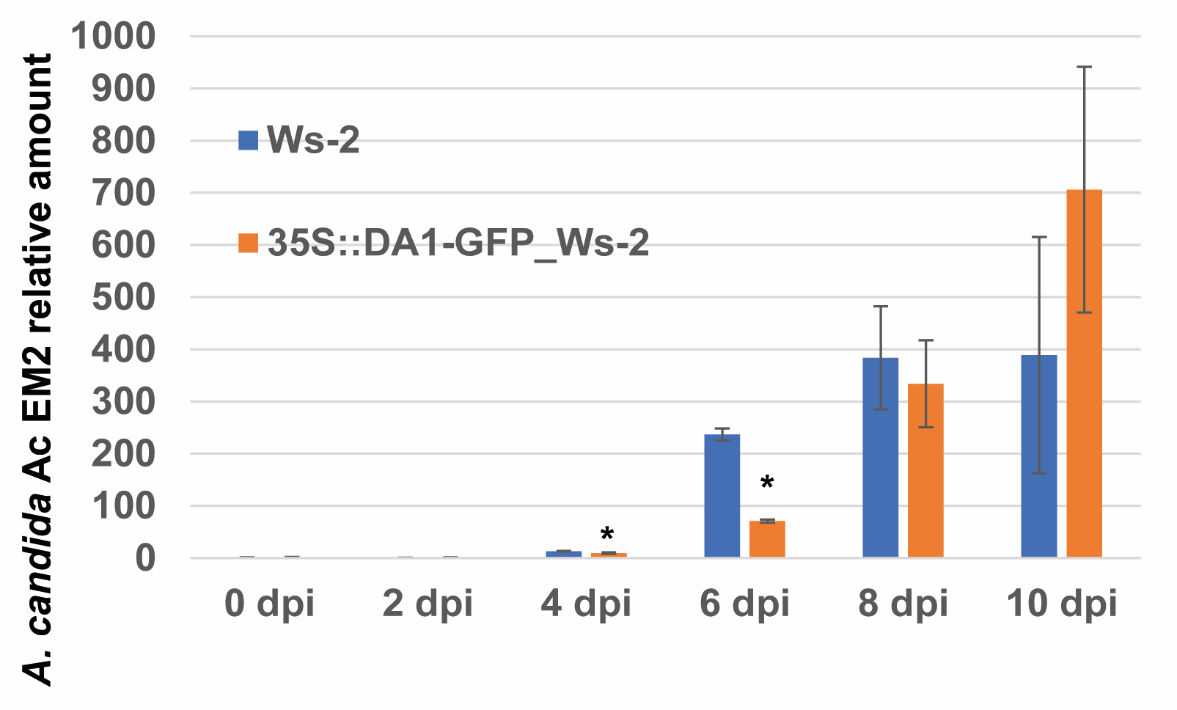

**Supplementary Figure 7. *Albugo candida* Em2 growth in Ws-2 leaves expressing *DA1-GFP* from the 35S promoter.**

Ws-2 leaves were inoculated at 0 days post-infection (dpi) with *A. candida* spores and growth was measured using q-PCR. Statistical significance was at *P < 0.05 based on a two-tailed Student’s t test. Error bars represent SD of the mean (Data in Supplementary Table 5).

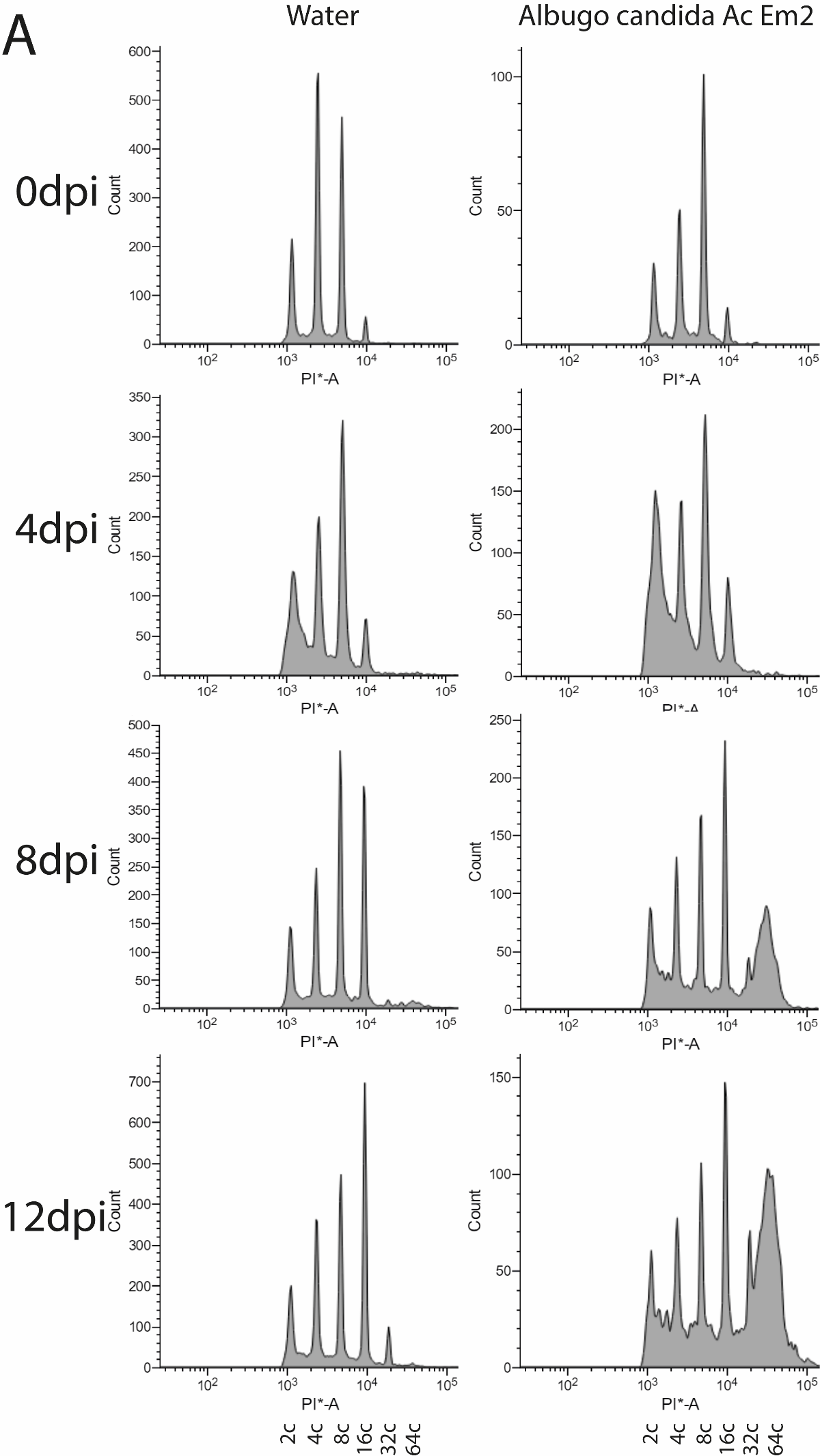

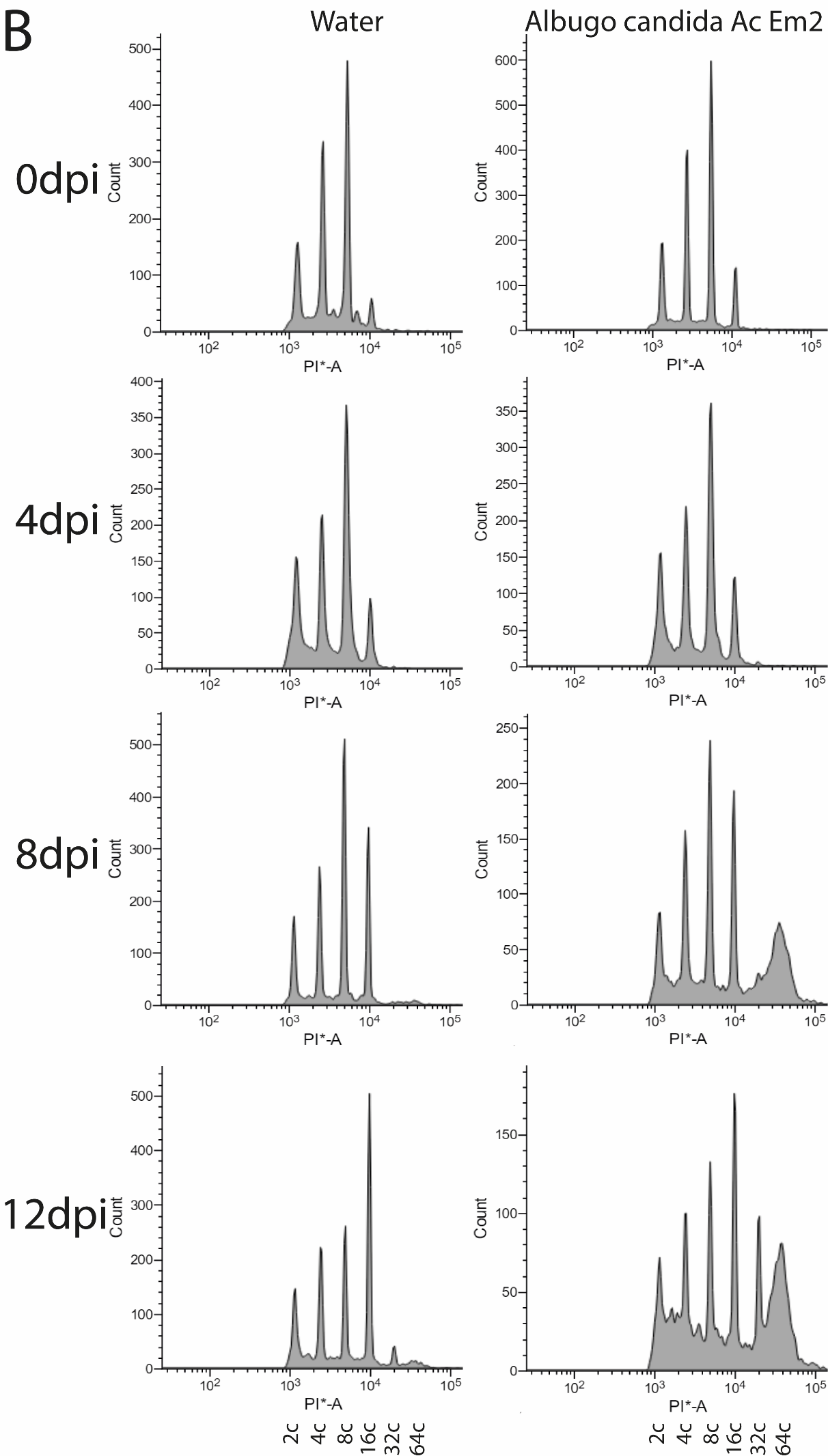

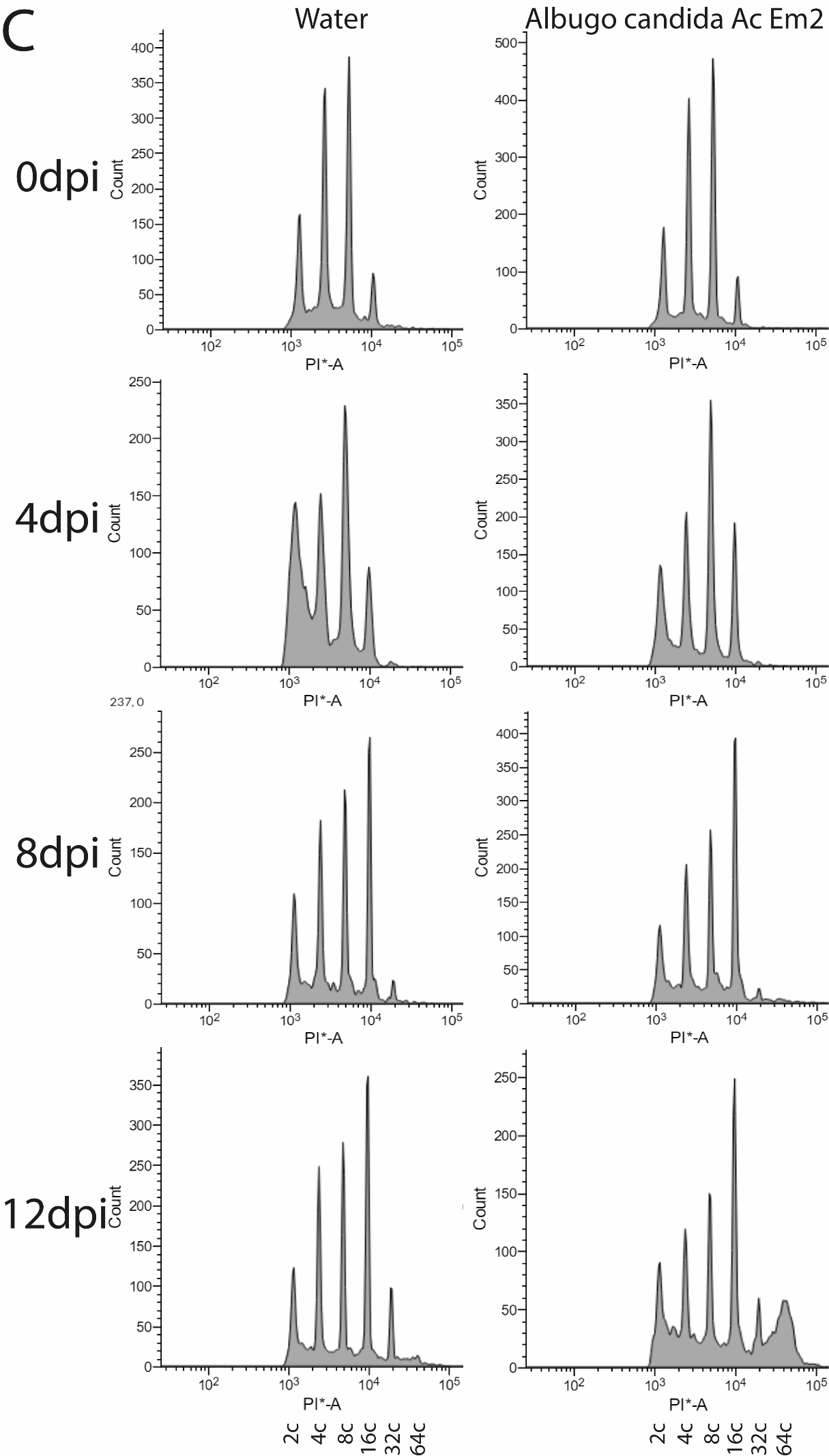

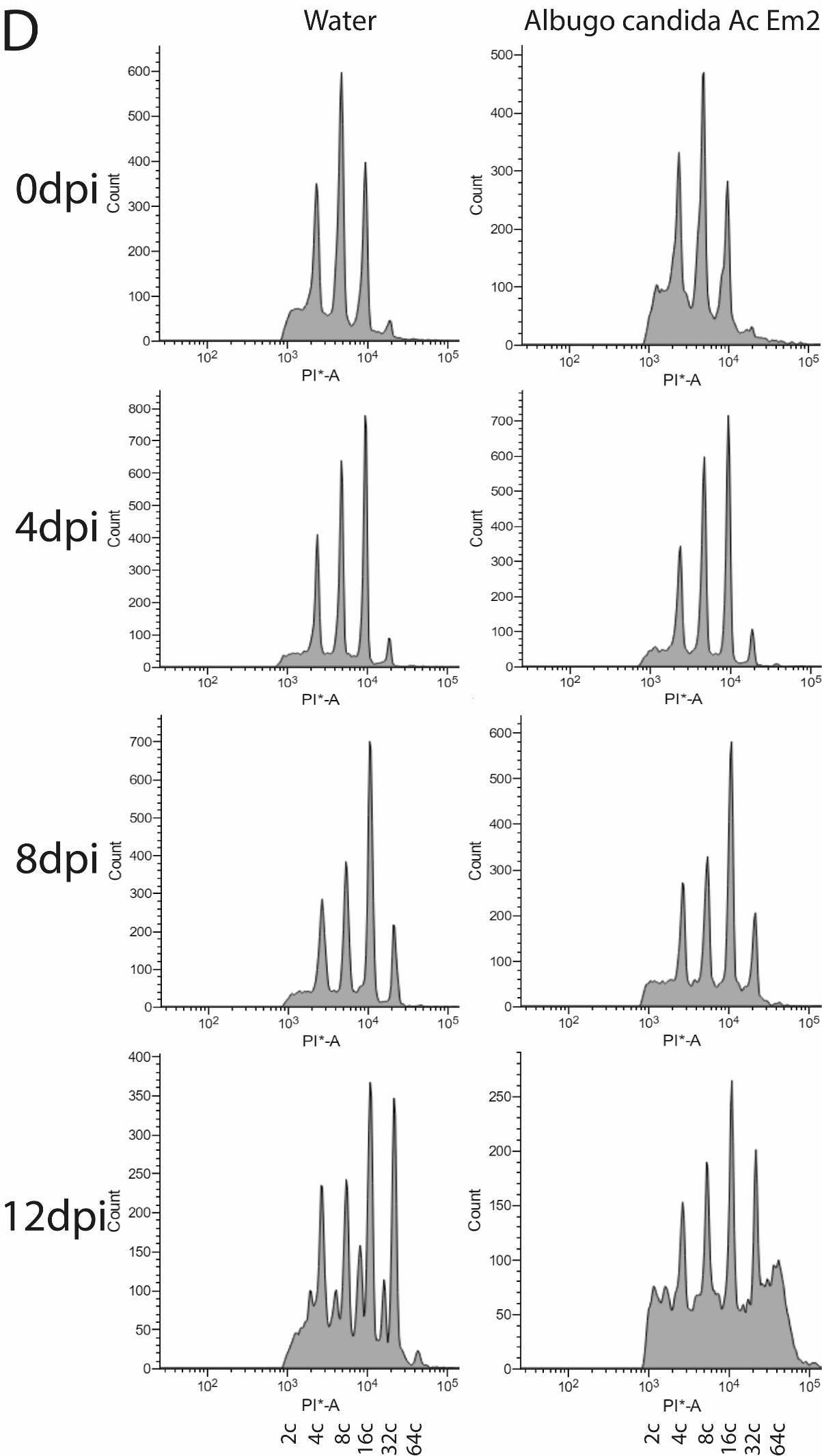

**Supplementary Figure 8. Leaf cell ploidy profiles of *Arabidopsis thaliana* accession Ws-2 lines infected by *Albugo candida* Ac Em2**

(A) Wild type Ws-2.

(B) BC7F2 line wt_42 with DA1 and DAR1 from Col-0 in Ws-2.

(C) BC7F2 line *da1-ko1dar1-1* with *da1-ko dar1-1* double mutant from Col-0 in Ws-2.

(D) *DA1* over-expression line 35S::-*DA1-GFP*_Ws-2

Water treatment was use as a control. Dpi: day post inoculation. 2c, 4c, 8c, 16c, 32c and 64c indicate the haploid copy numbers (Data in Supplementary Table 6).

**A**

```
DAR3 Crispr mutant protein sequences alignment
DAR3      MVRRKRQEEDEKIEIERVKEESLKLAKQAEEKRRLEESKEQGKRIQVDDDQLAKTTSKDK	60
dar3-2    MVRRKRQEEDEKIEIERVKEESLKLAKQAEEKRRLEESKEQGKRIQVDDDQLAKTTSKDK	60
dar3-3    MVRRKRQEEDEKIEIERVKEESLKLAKQAEEKRRLEESKEQGKRIQVDDDQLAKTTSKDK	60
```

************************************************************

**LIM domain**

```
DAR3      GQINHSKDVVEEDVNPPPSIDGKSEIGDGTSVNPRCLCCFHCHRPFVMHEILKKGKFHID	120
dar3-2    GQINHSKDVVEEDVNPPPSIDGKSEIGDGTSVNPRCLCCFHCHRPFVMHEILKKGKFHID	120
dar3-3    GQINHSKDVVEEDVNPPPSIDGKSEIGDGTSVNPRCLCCFHCHRPFVMHEILKKGKFHID	120
          ************************************************************
```

**LIM-like domain**

```
DAR3      CYKEYYRNRNCYVCQQKIPVNAEGIRKFSEHPFWKEKYCPIHDEDGTAKCCSCERLEPRG	180
dar3-2    CYKEYYRNRNCYVCQQKIPVNAEGIRKFSEHPFWKEKYCPIHDEDGTAKCCSCERLEPRG	180
dar3-3    CYKEYYRNRNCYVCQQKIPVNAEGIRKFSEHPFWKEKYCPIHDEDGTAKCCSCERLEPRG	180
          ************************************************************
```

```
DAR3      TNYVMLGDFRWLCIECMGSAVMDTNEVQPLHFEIREFFEGLFLKVDKEFALLLVEKQALN	240
dar3-2    TNYVMLGDFRWLCIECMGSAVMDTNEVQPLHFEIREFFEGLFLKVDKEFALLLVEKQALN	240
dar3-3    TNYVMLGDFRWLCIECMGSAVMDTNEVQPLHFEIREFFEGLFLKVDKEFALLLVEKQALN	240
          ************************************************************

DAR3      KAEEEEKIDYHRAAVTRGLCMSEEQIVPSIIKGPRMGPDNQLITDIVTESQRVSGFEVTG	300
dar3-2    KAEEEEKIDYHRAAVTRGLCMSEEQIVPSIIKGPRMGPVY*-------------------	280
dar3-3    KAEEEEKIDYHRAAVTRGLCMSEEQIVPSIIKGPRMGPDNQLITDIVTESQRVSGFEVTG	300
          **************************************
```

**peptidase active site**

```
DAR3      ILIIYGLPRLLTGYILAHEMMHAWLRLNGYKNLKLELEEGLCQALGLRWLESQTFASTDA	360
dar3-2    ------------------------------------------------------------	280
dar3-3    ILIIYGLPRLLTGYILAHEMMHAWLRLNGYKNLKLELEEGLCQALGLRWLESQTFASTDA	360


DAR3      AAAAAVASSSSFSSSTAPPAAITSKKSDDWSIFEKKLVEFCMNQIKEDDSPVYGLGFKQV	420
dar3-2    ------------------------------------------------------------	280
dar3-3    AAAAAVASSSSFSSDAAAAAVAAAVASSSSRYHVKEK*----------------------	397


DAR3      YEMMVSNNYNIKDTLKDIVSASNATPDSTV*				450
dar3-2    -------------------------------				280
dar3-3    -------------------------------				397
```

**B**

```
DAR3 Crispr mutant genomic sequences alignment
>GenomicDAR3        #1                  ATGGTACGTC GGAAAAGACA AGAAGAAGAT GAAAAGATTG
>DAR3_CDS           #1                  ATGGTACGTC GGAAAAGACA AGAAGAAGAT GAAAAGATTG
>dar3-2             #1                  ATGGTACGTC GGAAAAGACA AGAAGAAGAT GAAAAGATTG
>dar3-3             #1                  ATGGTACGTC GGAAAAGACA AGAAGAAGAT GAAAAGATTG
                                        ...........................................
                    #1                  ATGGTACGTC GGAAAAGACA AGAAGAAGAT GAAAAGATTG


>GenomicDAR3        #41                 AGATTGAAAG GGTTAAGGAA GAAAGTTTGA AGCTAGCTAA
>DAR3_CDS           #41                 AGATTGAAAG GGTTAAGGAA GAAAGTTTGA AGCTAGCTAA
>dar3-2             #41                 AGATTGAAAG GGTTAAGGAA GAAAGTTTGA AGCTAGCTAA
>dar3-3             #41                 AGATTGAAAG GGTTAAGGAA GAAAGTTTGA AGCTAGCTAA
                                        ...........................................
                    #41                 AGATTGAAAG GGTTAAGGAA GAAAGTTTGA AGCTAGCTAA


>GenomicDAR3        #81                 GCAGGAGGAA GAAAAGAGAA GATTAGAAGA GTCGAAGGAG
>DAR3_CDS           #81                 GCAGGCGGAA GAAAAGAGAA GATTAGAAGA GTCGAAGGAG
>dar3-2             #81                 GCAGGAGGAA GAAAAGAGAA GATTAGAAGA GTCGAAGGAG
>dar3-3             #81                 GCAGGAGGAA GAAAAGAGAA GATTAGAAGA GTCGAAGGAG
                                        ...........................................
                    #81                 GCAGGAGGAA GAAAAGAGAA GATTAGAAGA GTCGAAGGAG
                                             *

>GenomicDAR3        #121                CAAGGCAAAA GAATACAAGT GGACGATGAT CAGCTTGCAA
>DAR3_CDS           #121                CAAGGCAAAA GAATACAAGT GGACGATGAT CAGCTTGCAA
>dar3-2             #121                CAAGGCAAAA GAATACAAGT GGACGATGAT CAGCTTGCAA
>dar3-3             #121                CAAGGCAAAA GAATACAAGT GGACGATGAT CAGCTTGCAA
                                        ...........................................
                    #121                CAAGGCAAAA GAATACAAGT GGACGATGAT CAGCTTGCAA


>GenomicDAR3        #161                AGACTACTTC TAAAGATAAG GGACAAATTA ATCATTCCAA
>DAR3_CDS           #161                AGACTACTTC TAAAGATAAG GGACAAATTA ATCATTCCAA
>dar3-2             #161                AGACTACTTC TAAAGATAAG GGACAAATTA ATCATTCCAA
>dar3-3             #161                AGACTACTTC TAAAGATAAG GGACAAATTA ATCATTCCAA
                                        ...........................................
                    #161                AGACTACTTC TAAAGATAAG GGACAAATTA ATCATTCCAA


>GenomicDAR3        #201                AGATGTAGTG GAAGAAGATG TGAATCCTCC TCCTAGGTAA
>DAR3_CDS           #201                AGATGTAGTG GAAGAAGATG TGAATCCTCC TCCTAG::::
>dar3-2             #201                AGATGTAGTG GAAGAAGATG TGAATCCTCC TCCTAGGTAA
>dar3-3             #201                AGATGTAGTG GAAGAAGATG TGAATCCTCC TCCTAGGTAA
                                        ...........................................
                    #201                AGATGTAGTG GAAGAAGATG TGAATCCTCC TCCTAGGTAA


>GenomicDAR3        #241                AGCTATATCC TATATATATG TGTTCCACTG CGCAATTTTG
>DAR3_CDS           #241                :::::::::: :::::::::: :::::::::: ::::::::::
>dar3-2             #241                AGCTATATCC TATATATATG TGTTCCACTG CGCAATTTTG
>dar3-3             #241                AGCTATATCC TATATATATG TGTTCCACTG CGCAATTTTG
                                        ...........................................
                    #241                AGCTATATCC TATATATATG TGTTCCACTG CGCAATTTTG


>GenomicDAR3        #281                TATTTAGAGA TTTTTGGCTC AAATCAAGTA TTGTTTTACT
>DAR3_CDS           #281                :::::::::: :::::::::: :::::::::: ::::::::::
>dar3-2             #281                TATTTAGAGA TTTTTGGCTC AAATCAAGTA TTGTTTTACT
>dar3-3             #281                TATTTAGAGA TTTTTGGCTC AAATCAAGTA TTGTTTTACT
                                        ...........................................
                    #281                TATTTAGAGA TTTTTGGCTC AAATCAAGTA TTGTTTTACT


>GenomicDAR3        #321                TTACTTGTAG CAGTATTGAT GGCAAATCTG AGATTGGAGA
>DAR3_CDS           #321                :::::::::: :::TATTGAT GGCAAATCTG AGATTGGAGA
>dar3-2             #321                TTACTTGTAG CAGTATTGAT GGCAAATCTG AGATTGGAGA
>dar3-3             #321                TTACTTGTAG CAGTATTGAT GGCAAATCTG AGATTGGAGA
                                        ...........................................
                    #321                TTACTTGTAG CAGTATTGAT GGCAAATCTG AGATTGGAGA


>GenomicDAR3        #361                TGGAACATCT GTCAATCCTC GATGTAAATG TTGTTTTCAT
>DAR3_CDS           #361                TGGAACATCT GTCAATCCTC GATGTTTATG TTGTTTTCAT
>dar3-2             #361                TGGAACATCT GTCAATCCTC GATGTAAATG TTGTTTTCAT
>dar3-3             #361                TGGAACATCT GTCAATCCTC GATGTAAATG TTGTTTTCAT
                                        ...........................................
                    #361                TGGAACATCT GTCAATCCTC GATGTAAATG TTGTTTTCAT
                                                                   **

>GenomicDAR3        #401                TGCCACAGAC CATTTGTTAT GCACGAGGTC TAAAACCATA
>DAR3_CDS           #401                TGCCACAGAC CATTTGTTAT GCACGAG::: ::::::::::
>dar3-2             #401                TGCCACAGAC CATTTGTTAT GCACGAGGTC TAAAACCATA
>dar3-3             #401                TGCCACAGAC CATTTGTTAT GCACGAGGTC TAAAACCATA
                                        ...........................................
                    #401                TGCCACAGAC CATTTGTTAT GCACGAGGTC TAAAACCATA


>GenomicDAR3        #441                CATACAACTT GCAAACTCCT TTGTCTTGTT GTTTTCTGAT
>DAR3_CDS           #441                :::::::::: :::::::::: :::::::::: ::::::::::
>dar3-2             #441                CATACAACTT GCAAACTCCT TTGTCTTGTT GTTTTCTGAT
>dar3-3             #441                CATACAACTT GCAAACTCCT TTGTCTTGTT GTTTTCTGAT
                                        ...........................................
                    #441                CATACAACTT GCAAACTCCT TTGTCTTGTT GTTTTCTGAT


>GenomicDAR3        #481                GTATATAACT CCTTAGATTT TGAAGAAGGG AAAATTTCAC
>DAR3_CDS           #481                :::::::::: ::::::ATTT TGAAGAAGGG AAAATTTCAC
>dar3-2             #481                GTATATAACT CCTTAGATTT TGAAGAAGGG AAAATTTCAC
>dar3-3             #481                GTATATAACT CCTTAGATTT TGAAGAAGGG AAAATTTCAC
                                        ...........................................
                    #481                GTATATAACT CCTTAGATTT TGAAGAAGGG AAAATTTCAC


>GenomicDAR3        #521                ATAGATTGCT ACAAGGAGTA CTACCGTAAT CGCAACTGCT
>DAR3_CDS           #521                ATAGATTGCT ACAAGGAGTA CTACCGTAAT CGCAACTGCT
>dar3-2             #521                ATAGATTGCT ACAAGGAGTA CTACCGTAAT CGCAACTGCT
>dar3-3             #521                ATAGATTGCT ACAAGGAGTA CTACCGTAAT CGCAACTGCT
                                        ...........................................
                    #521                ATAGATTGCT ACAAGGAGTA CTACCGTAAT CGCAACTGCT


>GenomicDAR3        #561                ATGTTTGCCA ACAAAAGGTA AATAAATTTT CATTAATTAG
>DAR3_CDS           #561                ATGTTTGCCA ACAAAAG::: :::::::::: ::::::::::
>dar3-2             #561                ATGTTTGCCA ACAAAAGGTA AATAAATTTT CATTAATTAG
>dar3-3             #561                ATGTTTGCCA ACAAAAGGTA AATAAATTTT CATTAATTAG
                                        ...........................................
                    #561                ATGTTTGCCA ACAAAAGGTA AATAAATTTT CATTAATTAG


>GenomicDAR3        #601                TTTGGTTTTA CAGATTGTAG TAAAGTGAAC AGAATTGGTT
>DAR3_CDS           #601                :::::::::: :::::::::: :::::::::: ::::::::::
>dar3-2             #601                TTTGGTTTTA CAGATTGTAG TAAAGTGAAC AGAATTGGTT
>dar3-3             #601                TTTGGTTTTA CAGATTGTAG TAAAGTGAAC AGAATTGGTT
                                        ...........................................
                    #601                TTTGGTTTTA CAGATTGTAG TAAAGTGAAC AGAATTGGTT


>GenomicDAR3        #641                TATTTATGAC TTAATGAATG TGAAATACAA CTCTTTCAGA
>DAR3_CDS           #641                :::::::::: :::::::::: :::::::::: :::::::::A
>dar3-2             #641                TATTTATGAC TTAATGAATG TGAAATACAA CTCTTTCAGA
>dar3-3             #641                TATTTATGAC TTAATGAATG TGAAATACAA CTCTTTCAGA
                                        ...........................................
                    #641                TATTTATGAC TTAATGAATG TGAAATACAA CTCTTTCAGA


>GenomicDAR3        #681                TTCCTGTAAA TGCGGAAGGT ATAAGAAAGT TCAGTGAGCA
>DAR3_CDS           #681                TTCCTGTAAA TGCGGAAGGT ATAAGAAAGT TCAGTGAGCA
>dar3-2             #681                TTCCTGTAAA TGCGGAAGGT ATAAGAAAGT TCAGTGAGCA
>dar3-3             #681                TTCCTGTAAA TGCGGAAGGT ATAAGAAAGT TCAGTGAGCA
                                        ...........................................
                    #681                TTCCTGTAAA TGCGGAAGGT ATAAGAAAGT TCAGTGAGCA


>GenomicDAR3        #721                TCCGTTCTGG AAGGAGAAAT ACTGTCCTAT TCATGATGAG
>DAR3_CDS           #721                TCCGTTCTGG AAGGAGAAAT ACTGTCCTAT TCATGATGAG
>dar3-2             #721                TCCGTTCTGG AAGGAGAAAT ACTGTCCTAT TCATGATGAG
>dar3-3             #721                TCCGTTCTGG AAGGAGAAAT ACTGTCCTAT TCATGATGAG
                                        ...........................................
                    #721                TCCGTTCTGG AAGGAGAAAT ACTGTCCTAT TCATGATGAG


>GenomicDAR3        #761                GATGGAACTG CCAAGTGTTG CAGCTGTGAA AGATTAGAGG
>DAR3_CDS           #761                GATGGAACTG CCAAGTGTTG CAGCTGTGAA AGATTAGAG:
>dar3-2             #761                GATGGAACTG CCAAGTGTTG CAGCTGTGAA AGATTAGAGG
>dar3-3             #761                GATGGAACTG CCAAGTGTTG CAGCTGTGAA AGATTAGAGG
                                        ...........................................
                    #761                GATGGAACTG CCAAGTGTTG CAGCTGTGAA AGATTAGAGG


>GenomicDAR3        #801                TACTAATATA ATATTTGATC GAGTTTAGTT ATAAACAGAG
>DAR3_CDS           #801                :::::::::: :::::::::: :::::::::: ::::::::::
>dar3-2             #801                TACTAATATA ATATTTGATC GAGTTTAGTT ATAAACAGAG
>dar3-3             #801                TACTAATATA ATATTTGATC GAGTTTAGTT ATAAACAGAG
                                        ...........................................
                    #801                TACTAATATA ATATTTGATC GAGTTTAGTT ATAAACAGAG


>GenomicDAR3        #841                AAGAAAAAGC CTTGTGGCTT ATGACTTGTG ATTTGCAGCC
>DAR3_CDS           #841                :::::::::: :::::::::: :::::::::: ::::::::CC
>dar3-2             #841                AAGAAAAAGC CTTGTGGCTT ATGACTTGTG ATTTGCAGCC
>dar3-3             #841                AAGAAAAAGC CTTGTGGCTT ATGACTTGTG ATTTGCAGCC
                                        ...........................................
                    #841                AAGAAAAAGC CTTGTGGCTT ATGACTTGTG ATTTGCAGCC


>GenomicDAR3        #881                TAGGGGAACG AACTATGTAA TGCTTGGTGA TTTCCGGTGG
>DAR3_CDS           #881                TAGGGGAACG AACTATGTAA TGCTTGGTGA TTTCCGGTGG
>dar3-2             #881                TAGGGGAACG AACTATGTAA TGCTTGGTGA TTTCCGGTGG
>dar3-3             #881                TAGGGGAACG AACTATGTAA TGCTTGGTGA TTTCCGGTGG
                                        ...........................................
                    #881                TAGGGGAACG AACTATGTAA TGCTTGGTGA TTTCCGGTGG


>GenomicDAR3        #921                CTATGTATAG AGTGTATGGG ATCAGCGGTT ATGGATACTA
>DAR3_CDS           #921                CTATGTATAG AGTGTATGGG ATCAGCGGTT ATGGATACTA
>dar3-2             #921                CTATGTATAG AGTGTATGGG ATCAGCGGTT ATGGATACTA
>dar3-3             #921                CTATGTATAG AGTGTATGGG ATCAGCGGTT ATGGATACTA
                                        ...........................................
                    #921                CTATGTATAG AGTGTATGGG ATCAGCGGTT ATGGATACTA


>GenomicDAR3        #961                ACGAAGTCCA GCCTTTGCAC TTTGAAATCC GTGAATTCTT
>DAR3_CDS           #961                ACGAAGTCCA GCCTTTGCAC TTTGAAATCC GTGAATTCTT
>dar3-2             #961                ACGAAGTCCA GCCTTTGCAC TTTGAAATCC GTGAATTCTT
>dar3-3             #961                ACGAAGTCCA GCCTTTGCAC TTTGAAATCC GTGAATTCTT
                                        ...........................................
                    #961                ACGAAGTCCA GCCTTTGCAC TTTGAAATCC GTGAATTCTT


>GenomicDAR3        #1001               CGAAGGCTTG TTCTTGAAGG TTGACAAAGA ATTTGCTCTG
>DAR3_CDS           #1001               CGAAGGCTTG TTCTTGAAGG TTGACAAAGA ATTTGCTCTG
>dar3-2             #1001               CGAAGGCTTG TTCTTGAAGG TTGACAAAGA ATTTGCTCTG
>dar3-3             #1001               CGAAGGCTTG TTCTTGAAGG TTGACAAAGA ATTTGCTCTG
                                        ...........................................
                    #1001               CGAAGGCTTG TTCTTGAAGG TTGACAAAGA ATTTGCTCTG


>GenomicDAR3        #1041               CTTCTTGTCG AGAAACAAGC GCTTAATAAA GCTGAGGAAG
>DAR3_CDS           #1041               CTTCTTGTCG AGAAACAAGC GCTTAATAAA GCTGAGGAAG
>dar3-2             #1041               CTTCTTGTCG AGAAACAAGC GCTTAATAAA GCTGAGGAAG
>dar3-3             #1041               CTTCTTGTCG AGAAACAAGC GCTTAATAAA GCTGAGGAAG
                                        ...........................................
                    #1041               CTTCTTGTCG AGAAACAAGC GCTTAATAAA GCTGAGGAAG


>GenomicDAR3        #1081               AAGAGAAGAT CGTGAGTATA TTTAGAAAAC ACATCCCTCA
>DAR3_CDS           #1081               AAGAGAAGAT CG:::::::: :::::::::: ::::::::::
>dar3-2             #1081               AAGAGAAGAT CGTGAGTATA TTTAGAAAAC ACATCCCTCA
>dar3-3             #1081               AAGAGAAGAT CGTGAGTATA TTTAGAAAAC ACATCCCTCA
                                        ...........................................
                    #1081               AAGAGAAGAT CGTGAGTATA TTTAGAAAAC ACATCCCTCA


>GenomicDAR3        #1121               TCACTTTGTA ACTCATCTTT GTTTCTCATA TTTATGTCAA
>DAR3_CDS           #1121               :::::::::: :::::::::: :::::::::: ::::::::::
>dar3-2             #1121               TCACTTTGTA ACTCATCTTT GTTTCTCATA TTTATGTCAA
>dar3-3             #1121               TCACTTTGTA ACTCATCTTT GTTTCTCATA TTTATGTCAA
                                        ...........................................
                    #1121               TCACTTTGTA ACTCATCTTT GTTTCTCATA TTTATGTCAA


>GenomicDAR3        #1161               TACAAACAGG ACTACCATCG TGCAGCCGTA ACCAGAGGTC
>DAR3_CDS           #1161               :::::::::: ACTACCATCG TGCAGCCGTA ACCAGAGGTC
>dar3-2             #1161               TACAAACAGG ACTACCATCG TGCAGCCGTA ACCAGAGGTC
>dar3-3             #1161               TACAAACAGG ACTACCATCG TGCAGCCGTA ACCAGAGGTC
                                        ...........................................
                    #1161               TACAAACAGG ACTACCATCG TGCAGCCGTA ACCAGAGGTC


>GenomicDAR3        #1201               TTTGCATGTC TGAAGAGCAA ATTGTCCCAA GTGTAAGGCC
>DAR3_CDS           #1201               TTTGCATGTC TGAAGAGCAA ATTGTCCCAA GT::::::::
>dar3-2             #1201               TTTGCATGTC TGAAGAGCAA ATTGTCCCAA GTGTAAGGCC
>dar3-3             #1201               TTTGCATGTC TGAAGAGCAA ATTGTCCCAA GTGTAAGGCC
                                        ...........................................
                    #1201               TTTGCATGTC TGAAGAGCAA ATTGTCCCAA GTGTAAGGCC


>GenomicDAR3        #1241               AATCAATATA CACACTTTCA GAGACTAAGA GCAATAATGT
>DAR3_CDS           #1241               :::::::::: :::::::::: :::::::::: ::::::::::
>dar3-2             #1241               AATCAATATA CACACTTTCA GAGACTAAGA GCAATAATGT
>dar3-3             #1241               AATCAATATA CACACTTTCA GAGACTAAGA GCAATAATGT
                                        ...........................................
                    #1241               AATCAATATA CACACTTTCA GAGACTAAGA GCAATAATGT


>GenomicDAR3        #1281               CTAACTAGAT TTTCACTTTG GCTCCACTTC TATATATACA
>DAR3_CDS           #1281               :::::::::: :::::::::: :::::::::: ::::::::::
>dar3-2             #1281               CTAACTAGAT TTTCACTTTG GCTCCACTTC TATATATACA
>dar3-3             #1281               CTAACTAGAT TTTCACTTTG GCTCCACTTC TATATATACA
                                        ...........................................
                    #1281               CTAACTAGAT TTTCACTTTG GCTCCACTTC TATATATACA


>GenomicDAR3        #1321               GATAATAAAA GGGCCACGGA TGGGACCGGA CAATCAGCTA
>DAR3_CDS           #1321               :ATAATAAAA GGGCCACGGA TGGGACCGGA CAATCAGCTA
>dar3-2             #1321               GATAATAAAA GGGCCACGGA TGGGACCGGT CTA:CTGATG
>dar3-3             #1321               GATAATAAAA GGGCCACGGA TGGGACCGGA CAATCAGCTA
                                        ...........................................
                    #1321               GATAATAAAA GGGCCACGGA TGGGACCGGA CAATCAGCTA
                                                                       *  * * * * *

>GenomicDAR3        #1361               ATAACAGACA TAGTTACAGA GTCTCAAAGA GTGAGTGGAT
>DAR3_CDS           #1361               ATAACAGACA TAGTTACAGA GTCTCAAAGA GTGAGTGGAT
>dar3-2             #1361               CT:ACGGTC: TCG:AAAATA GA:::::::: ::::::::::
>dar3-3             #1361               ATAACAGACA TAGTTACAGA GTCTCAAAGA GTGAGTGGAT
                                        ...........................................
                    #1361               ATAACAGACA TAGTTACAGA GTCTCAAAGA GTGAGTGGAT
                                        * *  * * *  * ** * *   *

>GenomicDAR3        #1401               TCGAGGTTAC AGGGATTCTC ATCATATATG GACTTCCTAG
>DAR3_CDS           #1401               TCGAGGTTAC AGGGATTCTC ATCATATATG GACTTCCTAG
>dar3-2             #1401               :::::::::: :::::::::: :::::::::: ::::::::::
>dar3-3             #1401               TCGAGGTTAC AGGGATTCTC ATCATATATG GACTTCCTAG
                                        ...........................................
                    #1401               TCGAGGTTAC AGGGATTCTC ATCATATATG GACTTCCTAG


>GenomicDAR3        #1441               GTTACTAACA GGATATATCT TGGCTCACGA GATGATGCAT
>DAR3_CDS           #1441               GTTACTAACA GGATATATCT TGGCTCACGA GATGATGCAT
>dar3-2             #1441               :::::::::: :::::::::: :::::::::: ::::::::::
>dar3-3             #1441               GTTACTAACA GGATATATCT TGGCTCACGA GATGATGCAT
                                        ...........................................
                    #1441               GTTACTAACA GGATATATCT TGGCTCACGA GATGATGCAT


>GenomicDAR3        #1481               GCTTGGCTTA GACTCAATGG TACGTATTAA CTATGATTTC
>DAR3_CDS           #1481               GCTTGGCTTA GACTCAATGG T::::::::: ::::::::::
>dar3-2             #1481               :::::::::: :::::::::: :::::::::: ::::::::::
>dar3-3             #1481               GCTTGGCTTA GACTCAATGG TACGTATTAA CTATGATTTC
                                        ...........................................
                    #1481               GCTTGGCTTA GACTCAATGG TACGTATTAA CTATGATTTC


>GenomicDAR3        #1521               TCATCACCAA GAGACATGGA GAGAGTACCT CTTTTAATTT
>DAR3_CDS           #1521               :::::::::: :::::::::: :::::::::: ::::::::::
>dar3-2             #1521               :::::::::: :::::::::: :::::::::: ::::::::::
>dar3-3             #1521               TCATCACCAA GAGACATGGA GAGAGTACCT CTTTTAATTT
                                        ...........................................
                    #1521               TCATCACCAA GAGACATGGA GAGAGTACCT CTTTTAATTT


>GenomicDAR3        #1561               TTTTGTGATT TTGATCACAG GTTATAAGAA TCTTAAGTTA
>DAR3_CDS           #1561               :::::::::: :::::::::: ::TATAAGAA TCTTAAGTTA
>dar3-2             #1561               :::::::::: :::::::::: :::::::::: ::::::::::
>dar3-3             #1561               TTTTGTGATT TTGATCACAG GTTATAAGAA TCTTAAGTTA
                                        ...........................................
                    #1561               TTTTGTGATT TTGATCACAG GTTATAAGAA TCTTAAGTTA


>GenomicDAR3        #1601               GAGCTTGAAG AAGGATTATG CCAAGCGTTA GGTCTCAGAT
>DAR3_CDS           #1601               GAGCTTGAAG AAGGATTATG TCAAGCGTTA GGTCTCAGAT
>dar3-2             #1601               :::::::::: :::::::::: :::::::::: ::::::::::
>dar3-3             #1601               GAGCTTGAAG AAGGATTATG CCAAGCGTTA GGTCTCAGAT
                                        ...........................................
                    #1601               GAGCTTGAAG AAGGATTATG CCAAGCGTTA GGTCTCAGAT
                                                              *

>GenomicDAR3        #1641               GGTTGGAGTC TCTGACATTT GCATCTACTG ATGCTGCTGC
>DAR3_CDS           #1641               GGTTGGAGTC TCAGACATTT GCATCTACTG ATGCTGCTGC
>dar3-2             #1641               :::::::::: :::::::::: :::::::::: ::::::::::
>dar3-3             #1641               GGTTGGAGTC TCTGACATTT GCATCTACTG ATGCTGCTGC
                                        ...........................................
                    #1641               GGTTGGAGTC TCTGACATTT GCATCTACTG ATGCTGCTGC
                                                     *

>GenomicDAR3        #1681               TGCTGCTGCT GTAGCTTCTT CCTCTTCTTT TTCTTCTTCC
>DAR3_CDS           #1681               TGCTGCTGCT GTAGCTTCTT CCTCTTCTTT TTCTTCTTCC
>dar3-2             #1681               :::::::::: :::::::::: :::::::::: ::::::::::
>dar3-3             #1681               TGCTGCTGCT GTAGCTTCTT CCTCTTCTTT TTCTTCTGAT
                                        ...........................................
                    #1681               TGCTGCTGCT GTAGCTTCTT CCTCTTCTTT TTCTTCTTCC
                                                                                ***

>GenomicDAR3        #1721               ACTGC::::: :::::::::: :::::::::: ::::::TCCT
>DAR3_CDS           #1721               ACTGCT:::: :::::::::: :::::::::: :::::::CCT
>dar3-2             #1721               :::::::::: :::::::::: :::::::::: ::::::::::
>dar3-3             #1721               GCTGCTGCTG CTGCTGTAGC TGCTGCTGTA GCTTCTTCCT
                                        ...........................................
                    #1721               ACTGCTGCTG CTGCTGTAGC TGCTGCTGTA GCTTCTTCCT
                                        *

>GenomicDAR3        #1761               CCAGCCGCTA TCACGTCAAA GAAAAGTGAC GACTGGTCTA
>DAR3_CDS           #1761               CCAGCCGCTA TCACGTCAAA GAAAAGTGAC GACTGGTCTA
>dar3-2             #1761               CCAGCCGCTA TCACGTCAAA GAAAAGTGAC GACTGGTCTA
>dar3-3             #1761               CCAGCCGCTA TCACGTCAAA GAAAAGTGAC GACTGGTCTA
                                        ...........................................
                    #1761               CCAGCCGCTA TCACGTCAAA GAAAAGTGAC GACTGGTCTA


>GenomicDAR3        #1801               TTTTCGAGAA GAAGCTCGTA GAGTTTTGCA TGAATCAGAT
>DAR3_CDS           #1801               TTTTCGAGAA GAAGCTCGTA GAGTTTTGCA TGAATCAGAT
>dar3-2             #1801               TTTTCGAGAA GAAGCTCGTA GAGTTTTGCA TGAATCAGAT
>dar3-3             #1801               TTTTCGAGAA GAAGCTCGTA GAGTTTTGCA TGAATCAGAT
                                        ...........................................
                    #1801               TTTTCGAGAA GAAGCTCGTA GAGTTTTGCA TGAATCAGAT


>GenomicDAR3        #1841               AAAAGAGGAT GATTCACCGG TCTACGGTCT CGGATTCAAA
>DAR3_CDS           #1841               CAAAGAGGAT GATTCACCGG TCTACGGTCT CGGATTCAAA
>dar3-2             #1841               AAAAGAGGAT GATTCACCGG TCTACGGTCT CGGATTCAAA
>dar3-3             #1841               AAAAGAGGAT GATTCACCGG TCTACGGTCT CGGATTCAAA
                                        ...........................................
                    #1841               AAAAGAGGAT GATTCACCGG TCTACGGTCT CGGATTCAAA
                                        *

>GenomicDAR3        #1881               CAAGTTTACG AGATGATGGT CTCAAATAAC TACAACATTA
>DAR3_CDS           #1881               CAAGTTTACG AGATGATGGT CTCAAATAAC TACAACATTA
>dar3-2             #1881               CAAGTTTACG AGATGATGGT CTCAAATAAC TACAACATTA
>dar3-3             #1881               CAAGTTTACG AGATGATGGT CTCAAATAAC TACAACATTA
                                        ...........................................
                    #1881               CAAGTTTACG AGATGATGGT CTCAAATAAC TACAACATTA


>GenomicDAR3        #1921               AGGATACCCT TAAAGATATC GTTAGCGCTT CCAACGCTAC
>DAR3_CDS           #1921               AGGATACCCT TAAAGATATC GTTAGCGCTT CCAACGCTAC
>dar3-2             #1921               AGGATACCCT TAAAGATATC GTTAGCGCTT CCAACGCTAC
>dar3-3             #1921               AGGATACCCT TAAAGATATC GTTAGCGCTT CCAACGCTAC
                                        ...........................................
                    #1921               AGGATACCCT TAAAGATATC GTTAGCGCTT CCAACGCTAC


>GenomicDAR3        #1961               ACCAGATTCA ACGGTTTGA
>DAR3_CDS           #1961               ACCAGATTCA ACGGTTTGA
>dar3-2             #1961               ACCAGATTCA ACGGTTTGA
>dar3-3             #1961               ACCAGATTCA ACGGTTTGA
                                        ....................
                    #1961               ACCAGATTCA ACGGTTTGA
```

**Supplementary Figure 9. Designing the *dar3-2* and *dar3-3* CRISPR alleles of *DAR3* in the Ws-2 background.**

(A) Alignment of predicted protein sequence.

(B) Alignment of genomic sequence.

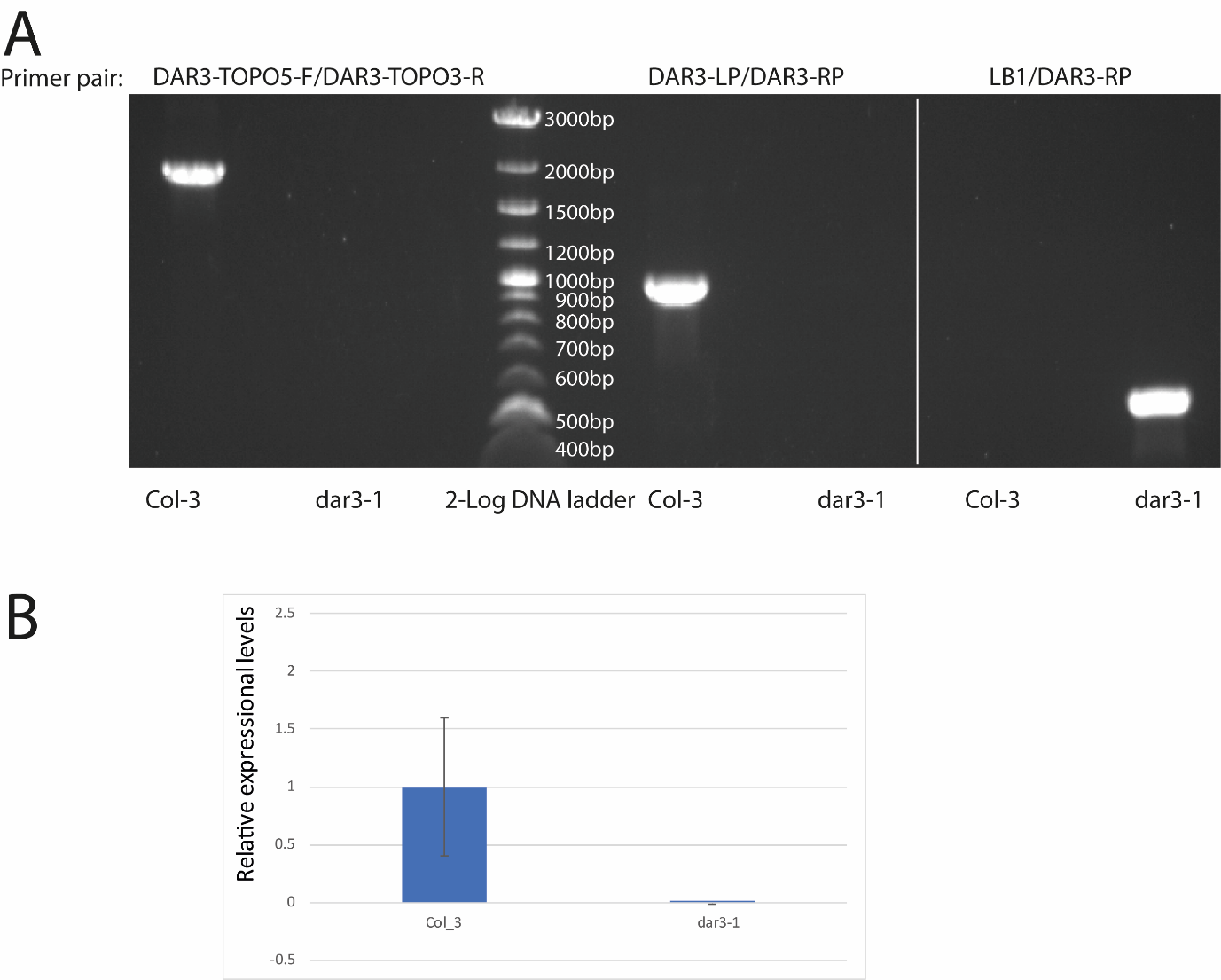

**Supplementary Figure 10. Characterising the *dar3-1* (SAIL_15_D09) T- DNA allele in the Col-3 background.**

(A) Genotyping the *dar3-1* mutant. Left: genomic DNA amplification of the whole DAR3 locus. Right: T-DNA insertion test. White bands indicate PCR products. Col-3 was used as a negative control.

(B) Relative expressional levels of wild type Col_3 and mutant *dar3-1*,determined by quantitative RT-PCR. Genomic DNA amplified by DAR3q-F/R and EF1-α_qF/R were used as internal references. Error bars represent SD of the mean (Supplementary Table 8).
